# Transcriptome analysis of heterogeneity in mouse model of metastatic breast cancer

**DOI:** 10.1101/2021.05.04.442511

**Authors:** Anastasia A. Ionkina, Gabriela Balderrama-Gutierrez, Kristian J. Ibanez, Steve Huy D. Phan, Angelique N. Cortez, Ali Mortazavi, Jennifer A. Prescher

## Abstract

Cancer metastasis is a complex process involving the spread of malignant cells from a primary tumor to distal organs. Understanding this cascade at a mechanistic level could provide critical new insights into the disease and potentially reveal new avenues for treatment. Transcriptome profiling of spontaneous cancer models is an attractive method to examine the dynamic changes accompanying tumor cell spread. However, such studies are often complicated by the underlying heterogeneity of the cell types involved. Here we performed a comprehensive analysis of tumor heterogeneity and organ tropism during breast cancer metastasis. We created a suite of organ-derived metastatic cell lines from the MMTV-PyMT mouse model. Bulk sequencing analyses uncovered tissue-specific genes across the different metastatic and primary tumor samples. We further investigated intratumoral heterogeneity by performing single-cell RNA-seq. These data revealed transcriptomes that may contribute to sub-clonal evolution during cancer progression. Collectively, the organ-derived cell lines provide a platform for comprehensive studies of metastatic breast cancer progression.

## Introduction

Despite recent advances in treatment and diagnosis, metastatic breast cancer remains a leading cause of death for women worldwide.^(Waks & Winer, 2019)^ Cancer metastasis is a complex process involving the spread of malignant cells from a primary tumor to distal organs.^(Vanharanta & Massague, 2013)(Steeg, 2016)^ Premalignant cells undergo dynamic cellular changes (i.e., epithelial to mesenchymal transition, EMT) to escape the primary tumor.^(Hanahan & Weinberg, 2011)^ These same cells undergo the reverse process (i.e., mesenchymal to epithelial transition, MET) to colonize metastatic sites.^(Harbeck et al., 2019)^ However, exactly which cells within a given primary tumor ultimately metastasize—and their final destinations—remains unclear.^(Hanahan & Weinberg, 2011; Harbeck et al., 2019; Montagner & Sahai, 2020)^

Transcriptome profiling of the dynamic cellular changes during tumorigenesis has the potential to improve our understanding of metastatic disease. Such analyses can reveal biomarkers associated with malignant progression. In one example, bulk RNA-sequencing (RNA-seq) revealed novel molecular pathways and differentially expressed genes (DEGs) associated with distinct stages of breast cancer progression.^(Cai et al., 2017)^ However, traditional profiling studies are complicated by the underlying heterogeneity of cancer progression. The contribution of distinct cell populations cannot be discerned using bulk RNA-seq alone.^(Zheng, Zhang, Zhao, Zhang, & Pollard, 2019)^ Single-cell RNA-seq (scRNA-seq) technology captures the complexity of cellular heterogeneity by mapping transcripts to individual cells.^(Olsen & Baryawno, 2018)^ This increase in cellular resolution facilitates the identification of additional molecular pathways and cell specific biomarkers. ^(H. Li et al., 2017; Yeo et al., 2020^)

Examining breast cancer remains challenging owing to a lack of models that capture cellular heterogeneity.^(Perlman, 2016)^ The surrounding microenvironment, cancer stem cells (CSC), and tumor dormancy all contribute to disease progression, and are difficult to replicate outside of living organisms. Suitable models must take into account the different tissue microenvironments that support cancer niches and resident cancer stem cells during metastatic progression.^(De Angelis, Francescangeli, La Torre, & Zeuner, 2019; Ghajar et al., 2013; Y. Li & Laterra, 2012; Takebe & Ivy, 2010)^ The MMTV-PyMT mouse model, in particular, is a well-established platform to study human breast cancer.^(Kersten, Salvagno, & de Visser, 2015; Ma et al., 2012)^ However, the variability in different tissue-dependent metastatic niches and the contribution of different cancer subclones to disease progression remains unclear. Transcriptome profiling of this cancer model could provide more insight into the mechanisms underlying dynamic changes in tumor progression.^(Liu, Dang, & Wang, 2018)^

We aimed to understand the transcriptome changes of organ-derived cancer cell isolates from MMTV-PyMT mice. Although metastatic progression from primary tumors to lung tissue is well studied in the MMTV-PyMT model, metastases to other distal organs and the significance of intratumor heterogeneity remain unclear.^(Gomez-Cuadrado, Tracey, Ma, Qian, & Brunton, 2017)^To gain insight, we established an array of metastatic cell lines harvested from MMTV-PyMT mice. Differential expression analyses were performed and used to examine the effects of cell heterogeneity on metastases and organ tropism. Correlations were found between CD44 expression and cellular growth markers across all metastatic cells. Data from scRNA-seq analyses further revealed tissue-specific gene expression patterns that mirror clinical data. Overall, the suite of clonal isolates provided a detailed depiction of cancer progression. The cell lines also establish a platform for future studies examining heterogeneity during metastatic disease and elucidating transcriptomic changes relevant to malignancy.

## Results

### Generation of breast cancer cell lines to examine tumor heterogeneity and metastatic disease

To gain insight into breast cancer heterogeneity, we derived a suite of tissue-specific metastatic cell lines from MMTV-PyMT mouse tumors (Fig. 1A). Tumors were harvested from the mammary fat pad (MFP) and tissues harboring distal metastases, including lymph nodes (LN), bone marrow (BM), and lungs (L). Samples were processed into single cell suspensions and further expanded. The organ-derived cultures were subjected to conditions that favored cancer cell outgrowth in vitro. Cells were ultimately sorted based on CD44 and EpCAM expression^(Marhaba et al., 2008)^ to remove fibroblasts from the samples. CD44 is routinely used as a marker of aggressive metastatic breast cancer. ^(Hen & Barkan, 2020)^ FACS sorting provided two populations: CD44^low^/EpCAM^high^ and CD44^high^/EpCAM^high^ (SI Fig. 1A-B). PCR was also used to confirm the presence of the PyMT viral antigen in the cell isolates (SI Fig. 1C). For the sorting and PCR assays, an established MMTV-PyMT cancer cell line (VO) and a common fibroblast cell line (3T3) were used as positive and negative controls, respectively. The tumorigenicity and metastatic propensity of the sorted MFP cell line was validated in vivo by injecting cultured cells into wild type female FVB mice (SI Fig. 1B). The presence of metastatic tumors was confirmed by harvesting LN, BM, and lung tissue from the re-injected mice. Single cell suspensions were formed and flow cytometry analysis confirmed the presence of CD44^low^/EpCAM^high^ and CD44^high^/EpCAM^high^ cells in the harvested tissues.

**Figure 1.**
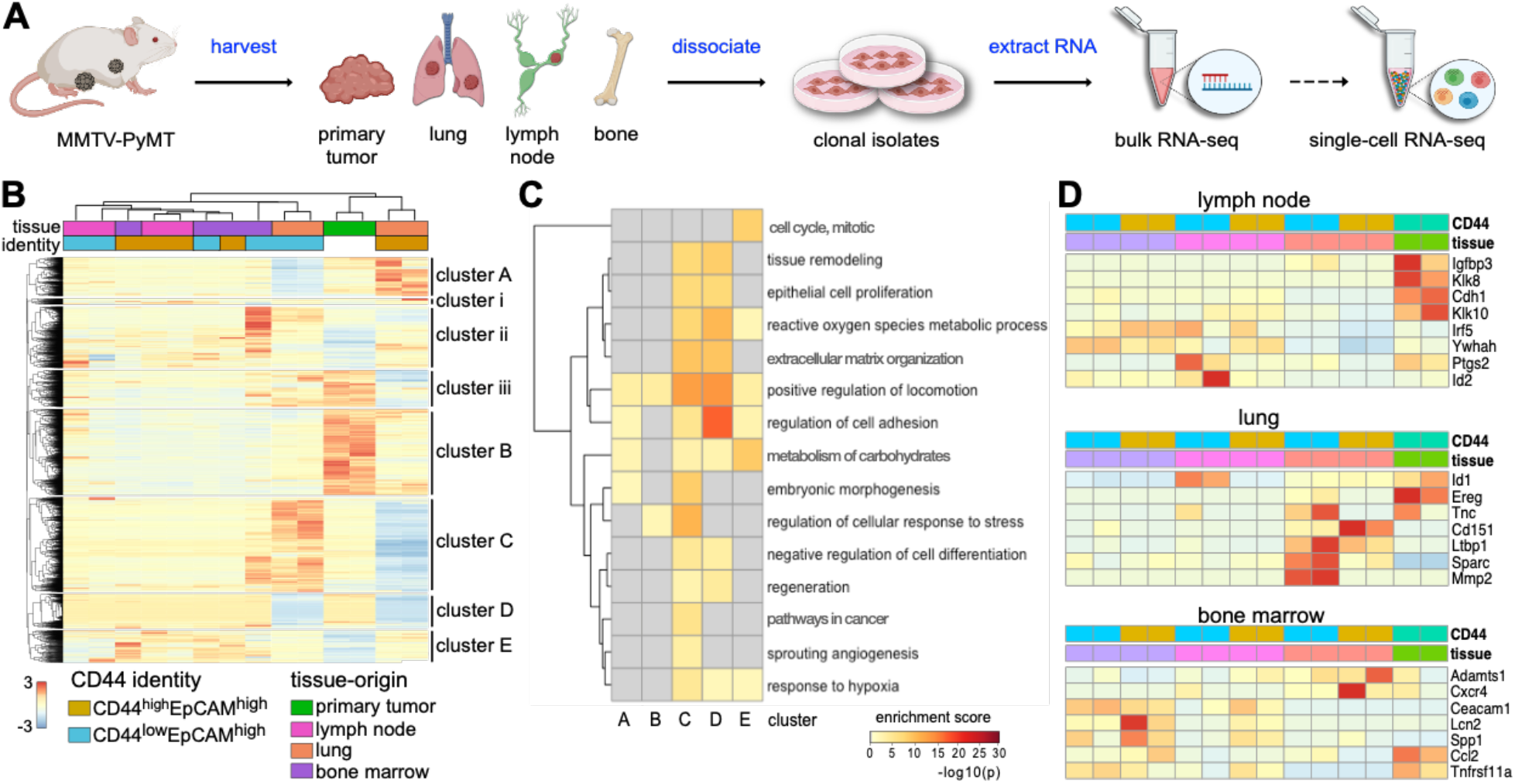
Clonal isolates from MMTV-PyMT breast cancer model exhibit distinct gene expression patterns. **(A)** Overview of cell isolation procedures and gene expression analyses. Tumors were harvested from mice and single cell suspensions were prepared. Cells were sorted based on CD44 and EpCAM expression. RNA was extracted for transcriptome profiling. Select samples were further analyzed via single-cell RNA-seq. **(B)** Heatmap of DEGs from tissue-specific metastatic cell lines and primary tumor sample. Expression levels for 5509 unique genes are shown. Values were normalized by row, and hierarchical clustering was used to sort the transcripts. Columns were clustered based on the tissue origin and CD44 expression level for each sample. Eight distinct gene clusters were observed, with clusters of interest annotated A-E. **(C)** GO-term enrichment analysis of clusters A-E from (B). GO terms were used to identify ontologies and biological processes relevant to cancer metastasis. Terms were also analyzed for signatures specific to the tissue of origin. The heat maps indicate the relative enrichment of the pathways across each cluster (columns). **(D)** Bulk RNA analysis revealed distinct gene expression patterns relevant to organ tropism. A panel of markers associated with tissue-tropic breast cancer metastases was examined across all samples. Clusters were assigned based on based on the tissue origin and CD44 expression level for each sample.

### Cancer cell lines exhibit distinct gene expression changes relative to metastatic progression

We used the tissue-derived cell lines to investigate transcriptional changes that occur during breast cancer metastasis. RNA was extracted from all cell samples and transcripts for established breast cancer genes were identified (SI Fig. 2).^(Cardoso et al., 2016; Institute, 2019; van ‘t Veer et al., 2002)^ Hierarchical clustering was performed on 5,509 DEGs. Eight distinct gene clusters (A-E; i-iii) were observed, as shown in Fig 1B. The transcripts were organized based on CD44 expression (CD44^high^/EpCAM^high^ or CD44^low^/EpCAM^high^) and tissue-origin (primary tumor, lymph node, lung or bone marrow). We focused on the five most prominent gene clusters (A-E) relevant to cancer progression for further analysis. Compared to the primary MFP tumor, the tissue-derived samples exhibited distinct upregulated and downregulated genes. Highly upregulated genes in MFP cells localized to cluster B. Lung-derived samples (CD44^low^ and CD44^high^) shared some similar transcriptomic changes (clusters D, E), but they also showed DEGs unique to their CD44 identity (clusters A, C). LN and BM samples trended similarly with MFP tumor cells, showing moderate expression of genes in cluster C.

To understand the biological relevance of the DEGs relevant to each cluster, we performed pathway enrichment analysis. Heat maps of the top 100 significant pathways revealed a multitude of cellular and molecular processes associated with cancer (SI Fig. 3, SI File 1). Pathways specifically relevant to cancer metastasis are shown in Fig. 1C, along with the corresponding enrichment score for each cluster. The upregulated genes for CD44^low^/EpCAM^high^ lung-derived cells in cluster C correlated with embryonic morphogenesis and hypoxia response pathways. Both of these pathways are critical to cancer cell growth in hostile environments. ^(Feng et al., 2018; Takebe, Warren, & Ivy, 2011)^ Some of the specific transcripts observed included those from the well-established cancer survival genes *ALDH1A1, SURVIVIN, XIAP, HSPG2, BCL9*, and *SOX4*. ^(Altieri, 2008; Evans et al., 2016)^ Interestingly, these same genes were downregulated in CD44^high^/EpCAM^high^ lung-derived cells (SI File 1). Cluster A was enriched in regulatory pathways associated with cell adhesion, correlating with the expression of *DDR1, HOXA7, MMP2, THBS1, TNFRSF14*, and *TGFB2*.^(Wasinski et al., 2020; Xu et al., 2005)^ Cluster A was also enriched in pathways associated with cellular locomotion, corroborated by the expression of *SERPINE1, PDGFA, ITGAV*, and *ITGB1BP1*.^(Heldin, 2013; Morandi et al., 2016)^ Both sets of upregulated genes were observed in CD44^high^/EpCAM^high^ lung-derived cells, but not MFP-derived or CD44^low^/EpCAM^high^ lung-derived cells. CD44^high^/EpCAM^high^ lung-derived cells also exhibited upregulated carbohydrate metabolism genes, pathways enriched in clusters A and E. Cluster E also correlated with other upregulated metabolic genes (*PFKFB3, SDC3*, and *GPC3*) in both lung-derived cell lines. These same genes were downregulated in the MFP cells.^(Atsumi et al., 2002; N. Li, Spetz, & Ho, 2020)^

Metastatic breast cancer cells are known to preferentially colonize specific organs, a process known as organotropism.^(Nguyen, Bos, & Massague, 2009)^ The cross talk between metastatic cells and the distal microenvironment leads to the formation of the pre-metastatic niche, which can influence cancer cell homing. ^(Minn et al., 2005; Obenauf & Massague, 2015)^ We examined whether the clonal isolates recapitulated features of organotropic metastases to lymph nodes, lung, and bone (Fig. 1D). Gratifyingly, we identified gene expression patterns that differed among the metastatic cell types based on their tissues of origin. LN-derived cells lines expressed genes relevant to metastatic lymphatic niches (e.g., *IRF5, YWHAH, PTGS2*, Fig. 1D) ^(Ross et al., 2020)^. MFP-associated genes *CDH1* and *IGFBP3* ^(W. Chen, Hoffmann, Liu, & Liu, 2018; Ross et al., 2020;)^ were also observed in the LN-derived lines, albeit to a lesser extent. In the case of the lung-derived cells, lung-tropic genes (e.g., *SPARC, MMP2, LTBP1, ID1*, and *CD151*) associated with high metastatic propensities ^(W. Chen, Hoffmann, Liu, & Liu, 2018; Ross et al., 2020;)^ were upregulated. *EREG*, a marker expressed by cells in the mammary gland,^(W. Chen, Hoffmann, Liu, & Liu, 2018)^ was downregulated in the lung-derived cells. BM-derived cells expressed genes associated with BM metastases, including *CEACAM1* and *LCN2*.^(Sun et al., 2019; Westbrook et al., 2016; Wood & Brown, 2020)^. Expression of *CCL2, ADAMST1* and *CXCR4* were also observed in BM-derived cells albeit to a lesser extent. Collectively, these transcriptome changes could contribute to sub-clonal evolution during cancer progression across the different metastatic niches.

We further compared the gene expression changes among the metastatic cells and to those from the primary tumor. Overall, we observed that samples derived from organs further away from the primary tumors had greater numbers of DEGs, regardless of the CD44 designation (SI File 2). Lung-derived samples exhibited the most DEGs (4,411 genes in total) compared to cells derived from the lymph node (1753 genes in total) or bone marrow (2,985 genes in total, SI Fig. 4). Volcano plots of DEGs from each tissue-derived metastatic cell line (CD44^high/low^/EpCAM^high^) compared to the primary tumor sample revealed genes involved in metastatic progression (SI Fig. 5 and SI File 2). Interestingly, some of the greatest differential expressions observed involved organotropism-associated genes (*MMP2* and *EREG*) identified in Fig. 1D.

We aimed to further characterize the metastatic cell lines via GO-term enrichment analysis. To this end, we examined gene expression changes relevant to metastatic progression, epithelial-to-mesenchymal transition (EMT), cellular proliferation and cell cycle control (Fig. 2). In the case of metastatic progression, we observed that MFP samples expressed high levels of classical markers associated with pre-metastatic lesions (e.g., *EREG, KRT14, CLDN7, KRT8, EMP1*, and *CLDN3*, Fig. 2A) ^(Minn et al., 2005; Obenauf & Massague, 2015)^. These markers were decreased in cells from distal metastatic sites (e.g., LN- and lung-derived cells). LN- and lung-derived cells, by contrast, exhibited upregulated levels of mesenchymal markers (e.g., *CCN5, ZEB1, VIM, SPARC* and *TGFB3*) ^(Ye et al., 2015)(Y. Li & Laterra, 2012; Sachidanandam et al., 2001)^. Expression levels were highest in CD44^high^/EpCAM^high^ lung-derived cells. Lung-derived cell lines also showed increased expression of well-established EMT markers (*SNAI1/2* and *CD63*)^(Ye et al., 2015)^ and markers associated with poor prognosis in patients (*FOXC1, AEBP1, SDC4*, and *IDH1*, Fig. 2B).^(W. Chen et al., 2018; Weigelt, Peterse, & van ‘t Veer, 2005)^ Interestingly, *SNAI1/2* and *CD63* expression were highest in CD44^low^/EpCAM^high^ lung-derived cells, while the poor prognosis indicators listed were higher in CD44^high^/EpCAM^high^ lung-derived cells. The upregulation of mesenchymal markers and downregulation of epithelial markers in lung-derived cells is indicative of cellular de-differentiation,^(Weigelt, Peterse, & van ‘t Veer, 2005) (Quail & Joyce, 2013)^ suggesting that the lung-derived cells recapitulate EMT.

**Figure 2.**
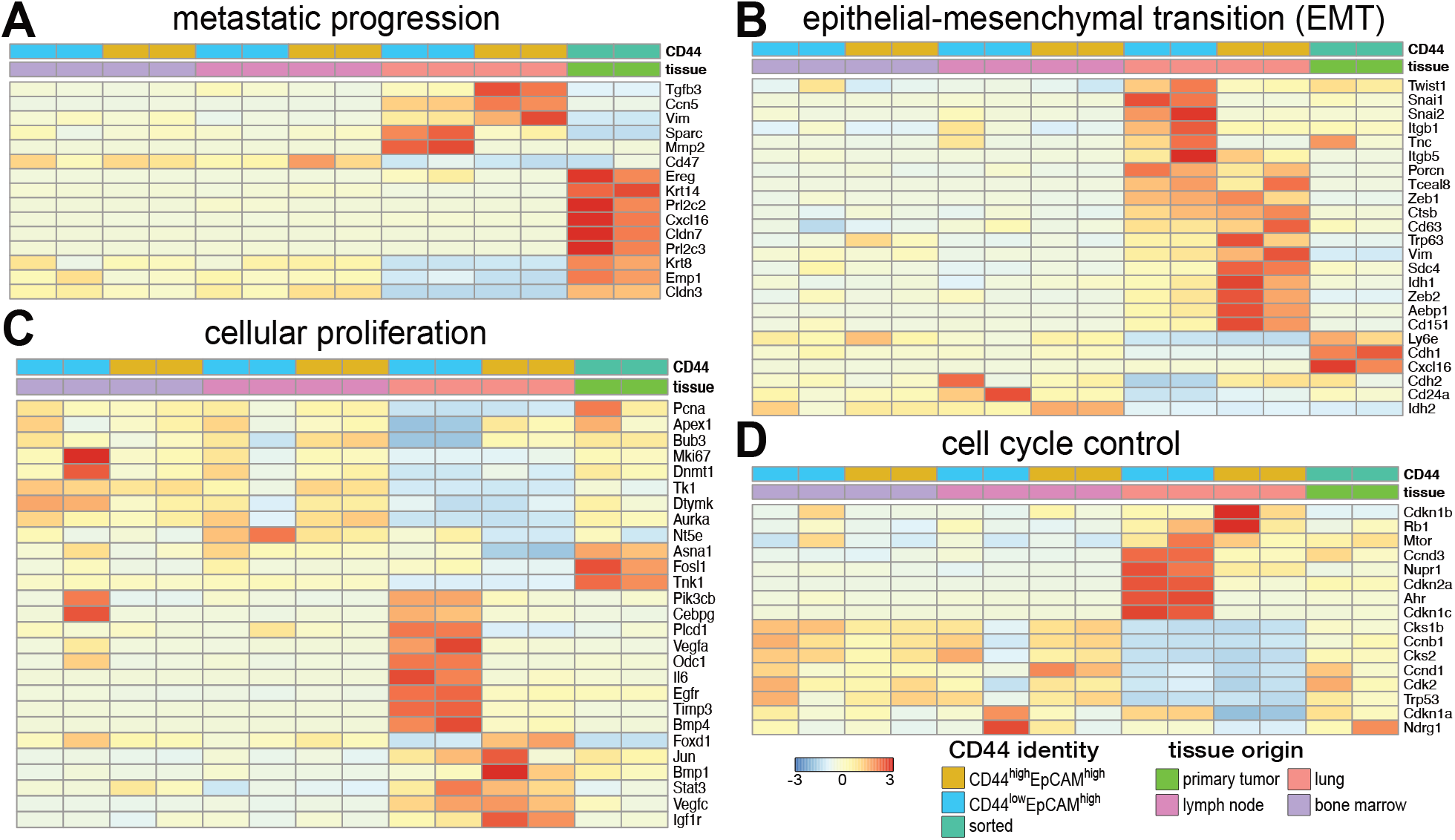
Cancer cell lines exhibit distinct gene expression patterns relative to metastatic disease progression. Bulk RNA analysis revealed differential gene expression patterns relevant to **(A)** metastatic progression, **(B)** epithelial-mesenchymal transition (EMT), **(C)** cellular proliferation, and **(D)** cell cycle control among the tissue-derived isolates and primary tumor. Columns were clustered based on CD44 expression and tissue origin as indicated. Select genes relevant to metastatic progression are displayed in the heat map.

EMT typically correlates with changes in cell proliferation and dysregulation of cell cycle control during cancer progression. These trends were apparent in the gene expression profiles for both CD44^high^ and CD44^low^ cells (Fig. 2C-D). As expected, the highly proliferative lung-derived CD44^high^ cells expressed low levels of growth arrest genes *CDKN1A* and *CDKN2A* (Fig 2D).^(Feng et al., 2018; Takebe, Warren, & Ivy, 2011)^ Interestingly, we observed a stark difference in gene expression for lung-derived CD44^low^ cells. Although these cells expressed genes relevant to cellular proliferation and angiogenesis, they exhibited upregulated levels of the growth arrest genes (Fig. 2D). Growth arrest signals could dampen the expression of other genes that are master regulators of downstream cell function. One such gene, mTOR, was expressed in the lung-derived CD44^low^ cells. The levels of mTOR were comparable to expression in CD44^high^ cells. From these results we postulate that lung CD44^low^ cells, albeit capable of cell division, are not dividing as rapidly as their CD44^high^ counterpart.

### Analyses of common biological pathways reveals intratumor heterogeneity

MMTV-PyMT has recently been used as a model to study the impacts of CD44 on metastases.^(C. Yang et al., 2019)^ Ex vivo analysis of tumors from a single micro-metastatic site revealed two subgroup of cells with differential CD44 expression. CD44 expression correlated with altered gene expression relevant to EMT and MET and differential growth rates.^(W. Chen et al., 2018; Weigelt, Peterse, & van ‘t Veer, 2005)^ Post-metastatic colonization, CD44 expression levels did not remain constant and were frequently switched between subgroups. The fluid transition from EMT to MET phenotypes demonstrates how complex and context-dependent breast cancer can be. The morphological changes induced by CD44 expression also affected the tumor-initiating capabilities of the tumor cells. However, the critical regenerative CSC populations were found in both CD44-expressing groups, warranting further characterization. Similar spectrums of behavior have been documented in other studies.^(Liu et al., 2018) (C. Yang et al., 2019) (K. Chen & Fraley, 2020)^

To more closely examine the impacts of CD44 expression in our organ-derived cells, we analyzed DEGs based on CD44 levels (SI File 3). We identified upregulated genes for CD44^low^/EpCAM^high^ (374 genes) and CD44^high^/EpCAM^high^ (276 genes) signatures across the suite of cells (SI Fig. 4). Some genes were shared across the tissue types. We also examined the GO terms and pathways associated with the DEGs from CD44^low^/EpCAM^high^ and CD44^high^/EpCAM^high^ samples. Clear differences were observed between the two CD44 signatures across the tissue-derived samples (Fig. 3A and SI File 4). Cells with CD44^high^ signatures exhibited an increase in GO terms and associated genes related to cellular proliferation, tumor aggression, and EMT. In contrast, CD44^low^ cells exhibited higher gene expression levels relevant to tumor microenvironment remodeling and stem cell markers. Interestingly, CD44^low^ signatures also correlated with signaling pathways known to be important for stem cell maintenance and Wnt-activated receptor activity. CD44^low^ signatures were further negatively correlated with cell differentiation pathways, supporting the idea of retained cellular dedifferentiation. The thrombospondin complex pathway, key in the maintenance of cancer stem cell dormancy in breast cancer, was also present in the CD44^low^ signature.^(Ghajar et al., 2013; Huang, Sun, Yuan, & Qiu, 2017; Quail & Joyce, 2013)^

**Figure 3.**
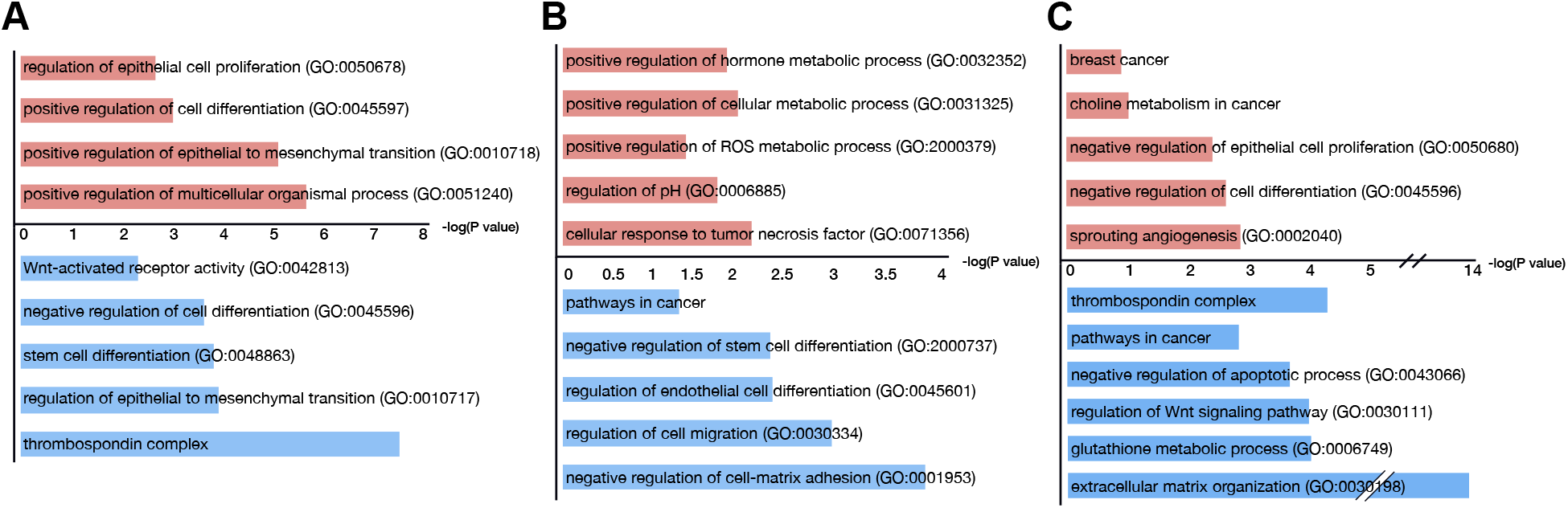
CD44 expression correlates with different transcriptome patterns in organ-derived cell lines. Differentially expressed genes for **(A)** CD44^high^ (red) and CD44^low^ (blue) EpCAM^high^ cells (across all samples) were used to generate GO terms. CD44^low^ cells exhibited high levels of expression for genes relevant to tumor microenvironment remodeling and tumor dormancy markers. CD44^high^ cells exhibited higher levels of gene expression associated with GO terms related to cellular proliferation, tumor aggression, and EMT. Differentially expressed genes and associated GO terms for CD44^high/low^ EpCAM^high^ cells from (B) lymph nodes and (C) lungs are also shown.

We further examined the intratumor heterogeneity of CD44^high^ versus CD44^low^ expression within single tumors. Volcano plots revealed DEGs in CD44^high^/EpCAM^high^ versus CD44^low^/EpCAM^high^ from lymph node-derived, lung-derived, and bone marrow-derived metastatic clonal isolates (SI Fig. 6A-C). The DEGs for these samples were also subjected to pathway enrichment analysis (Fig. 3B-C, SI Files 5-6). DEGs upregulated in lymph node-derived CD44^high^/EpCAM^high^ cells correlated with cellular metabolism and pH regulation (Fig. 3B) observed in aggressive cancer phenotypes. Similar pathways were not observed in the corresponding CD44^low^/EpCAM^high^ lymph node-derived cells. The DEGs for these cells, by contrast, were enriched in pathways regulating stem cell differentiation, cellular migration and cell-matrix adhesion (Fig. 3B). For the lung-derived samples, the CD44^high^/EpCAM^high^ cells exhibited upregulated DEGS relevant to cancer metabolism, cellular proliferation and angiogenesis (Fig. 3C). The DEGs for the corresponding CD44^low^/ EpCAM^high^ lung-derived cells were also enriched for pathways relevant to cancer progression and cellular metabolism, in addition to the thrombospondin complex and extracellular communication (Fig. 3C). Similar analyses were performed with bone marrow-derived cells to reveal unique GO terms and transcriptomic changes (SI Fig. 6D-E). Upregulated DEGs for BM-derived CD44^low^/ EpCAM^high^ cells were mainly associated with immune and cytokine activity.

Based on our DEG analyses, we examined additional known markers of breast cancer metabolism and extracellular remodeling across the entire set of organ-derived cell lines.^(Iliopoulos, Hirsch, Wang, & Struhl, 2011; Plaks, Kong, & Werb, 2015)^ We observed increased levels of cellular metabolism markers (e.g., *PLCB4, IGFBP7, IGFBP4, SHC2, PGRAMC1, MTHFD2*) specific to lung-derived CD44^low^ cells (Fig. 4A). Lung-derived CD44^high^ cells exhibited higher levels of *MPC1, MPC2, POGLUT1*, and *LARGE1* expression. Interestingly, we did not observe upregulation of other cancer-related drivers of energy consumption in either of the lung-derived cell lines compared to MFP-derived samples.^(Yi et al., 2020)^ Extracellular remodeling has been shown to improve cancer colonization and perpetuate dedifferentiated stem-like cellular states.^(Nagai, Ishikawa, Minami, & Nishita, 2020; Quail & Joyce, 2013)^ We identified markers relevant to extracellular remodeling known to promote the survival of metastatic lesions (*CD36, CD274, FOXC1*) in the lung-derived metastatic cells (Fig. 4B). The expression levels were noticeably enhanced in these samples compared to the MFP tumor. Lung-derived CD44^low^ cells also exhibited upregulated levels of genes associated with mesenchymal cells and more dedifferentiated phenotypes in advanced cancers (e.g., *FN1* and *POFUT2*).^(Y. Li & Laterra, 2012)^

**Figure 4.**
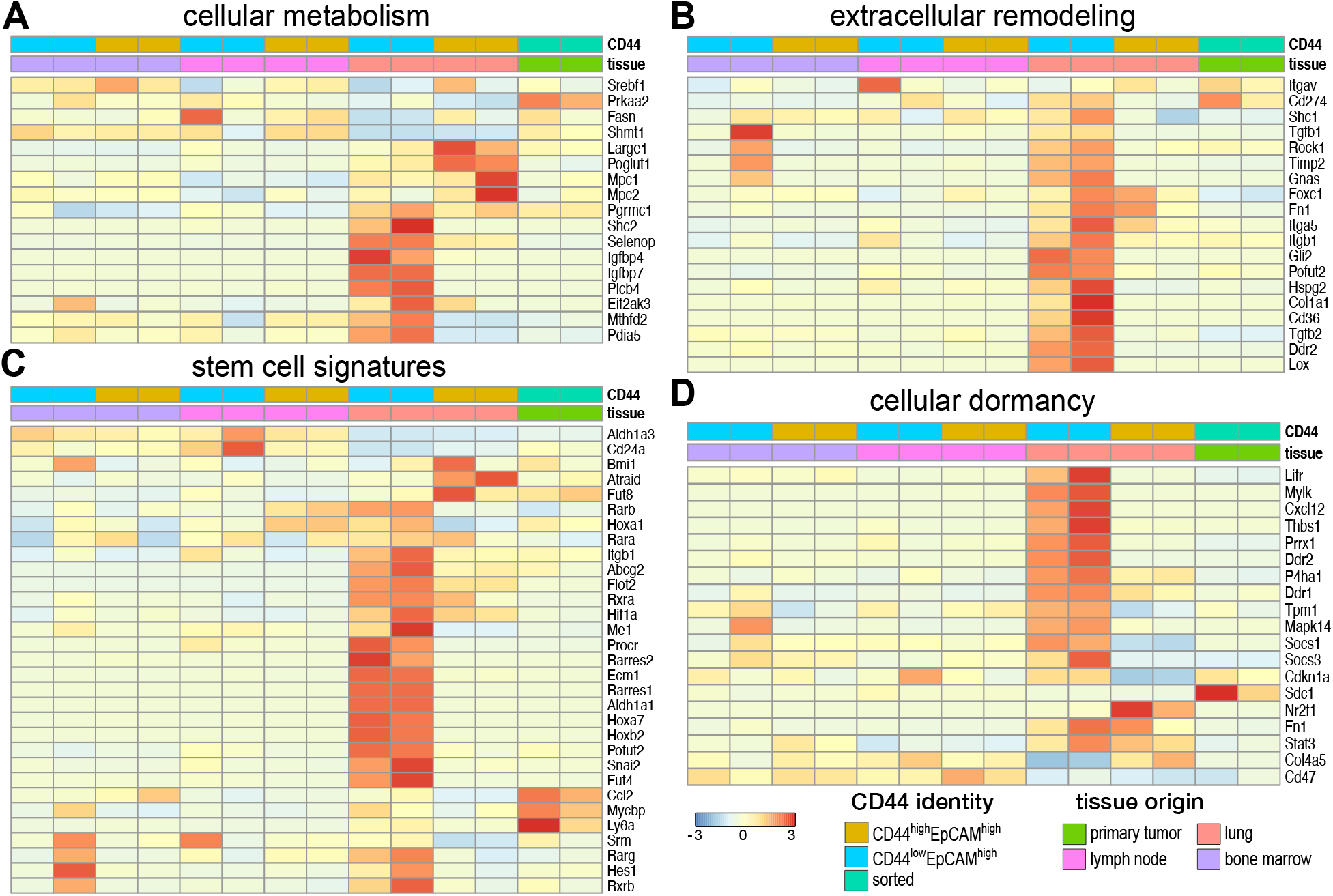
Intratumoral heterogeneity observed across organ-derived cell lines. Bulk RNA analysis of the organ-derived cell lines revealed distinct expression patterns relevant to **(A)** cellular metabolism, **(B)** extracellular remodeling of the microenvironment, **(C)** stem cell signatures, and **(D**) cellular dormancy among the cell lines. Clusters were assigned based on CD44 expression and the metastatic origin of each cell line as shown in the lower right.

### CD44^low^ cell lines exhibit classic signatures of stem cells

As noted earlier, the lung-derived CD44^low^ cells exhibited reduced levels of some markers of cellular proliferation and division (Fig 2C,D). These cells also expressed genes known to be important for CSC survival and function (Fig. 4B).^(L. Yang et al., 2020)^ Changes in gene expression relevant to matrix remodeling have been known to sustain CSCs in a functional, but dormant non-dividing state. To further examine whether the lung-derived CD44^low^ cells harbored CSC properties, we evaluated a panel of known breast cancer stem cell markers across our suite of metastatic isolates. Comparisons were made to the known cellular differentiation marker CD24 elevated ^(Vassalli, 2019)^. As shown in Fig. 4C, lung-derived CD44^low^/EpCAM^high^ cells exhibited an increase in the stem cell-associated markers and a decrease in CD24 expression. Furthermore, expression of *ALDH1A1*, a breast cancer-specific stem cell marker associated with resistance to some chemotherapies, was elevated.^(Tomita, Tanaka, Tanaka, & Hara, 2016; Vassalli, 2019)^ CSCs can endure some drug treatments and survive in metastatic environments due, in part, to their ability to modulate their metabolism and compensate for oxidative stress. ^(L. Yang et al., 2020)(Tomita et al., 2016)^ The lung-derived CD44^low^ cells exhibited gene expression profiles consistent with these phenotypes, including the upregulation of retinoic acid (RA) pathway (*RARA/B/G, RXRA/B, RARRES1/2*) essential for cell survival. Opposite trends were observed for lung-derived CD44^high^ cells. These cells expressed higher levels of genes associated with cell growth and aggressive metastases.

We further examined the entire suite of cells lines, for markers of breast cancer dormancy (Fig. 4D). The lung-derived CD44^low^ cells expressed higher levels of genes associated with the thrombospondin complex, a well-known dormancy marker in breast cancer.^(Barney et al., 2020; Ghajar et al., 2013)^ Additional markers, including *MAPK14, DDR1/2*, and *MYLK* were also observed among this cell population. Collectively, these data suggest that lung-derived CD44^low^ cells express CSC-relevant genes that can maintain cells in a dormant or low proliferative state. CSC identification remains challenging, though, owing to difficulties in isolation.^(Crabtree & Miele, 2018; Fico, Bousquenaud, Ruegg, & Santamaria-Martinez, 2019; Lobba, Forni, Carreira, & Sogayar, 2012)^

### Single cell RNA-seq reveals distinct clusters relevant to metastatic progression and intratumor heterogeneity

To further validate that our isolated cell lines capture the intratumoral heterogeneity observed during *de novo* disease progression, we performed scRNA-seq on a subset of metastatic samples. Based on the differential gene expression observed in bulk RNA-seq, we chose lung-derived CD44^low^ (lungL), lung-derived CD44^high^ (lungH), and LN-derived CD44^high^ (lymphH) cells for the analyses. In all, 4124 cells were used in the clustering analyses: lungL (334 cells), lungH (1085 cells), lymphH (1694 cells). Clusters were visualized using UMAP (Fig. 5A). Clusters for each of the three cell types—lungL (green), lungH (pink), and lymphH (blue)—were identified.

**Figure 5.**
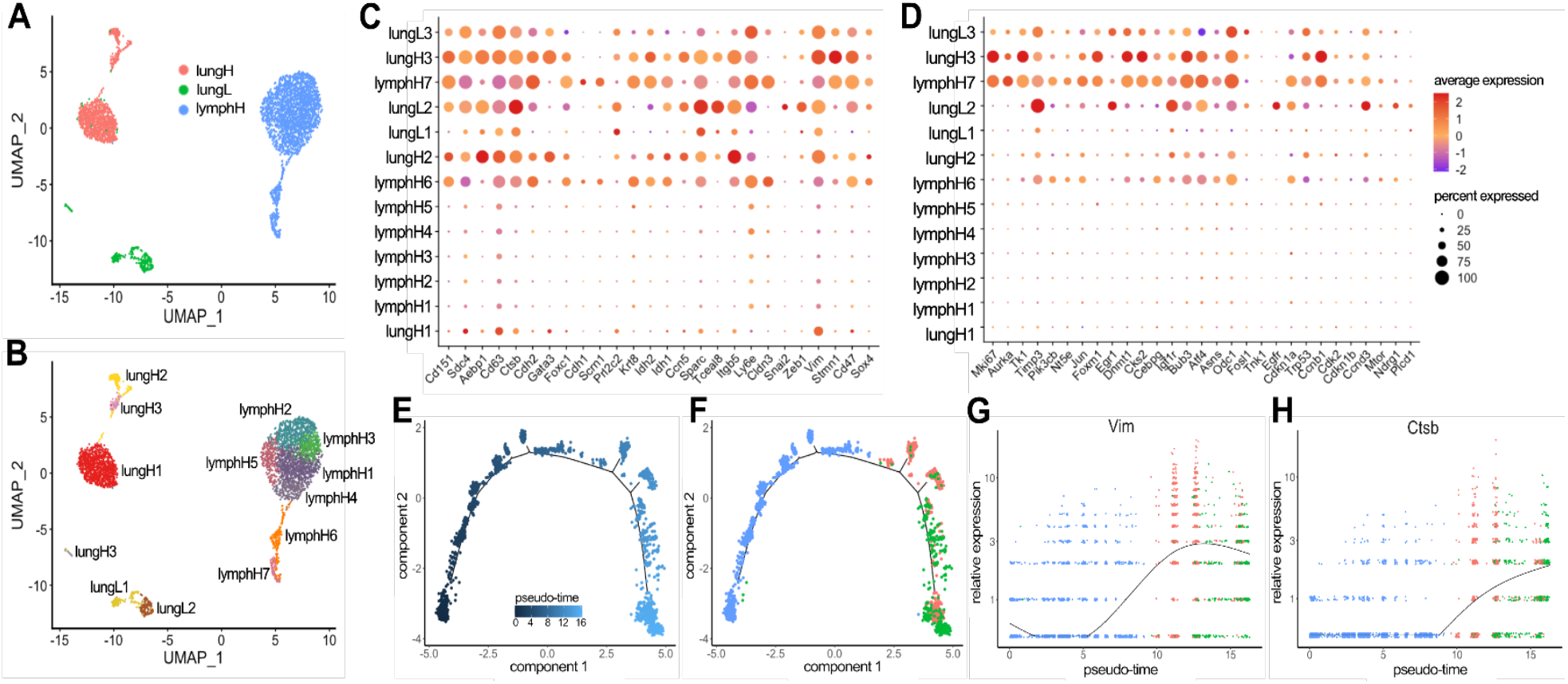
Single-cell RNA-seq revealed tissue-specific clusters and heterogeneity during metastatic disease progression. **(A)** Clustering of 4,124 cells that passed filtering. Tissue-specific clustering was observed for lungH, lungL, and lymphH cells. **(B)** Thirteen clusters were recovered based on a combination of tissue origin and CD44 signature. **(C, D)** Gene expression profiles for select cancer relevant genes (columns) relevant to **(C)** EMT and **(D)** cell proliferation among the thirteen clusters (rows). The size of each dot represents the percentage a specific gene is expressed compared to all other transcripts. The color gradient of the dot indicates the average expression of the gene. **(E)** Monocle2 pseudo-time analysis was performed and the metastatic trajectory of distinct cell clusters are shown. The color gradient indicates pseudo-time progression. **(F)** Cell tissues of origin indicated on the pseudo-time map (as in **A**). Changes in EMT-relevant genes from C were probed across pseudo-time and tissue of origin. **(G)** The expression of *VIM*, an EMT marker, increased over lymph node populations with maximum expression in lung high populations. This analysis was repeated for **(F)** *CTSB*, a marker of invasion, where the low expression across lymph node populations increased in expression across lung high and lung low populations.

We further examined the heterogeneity of the samples using hierarchical clustering (Fig. S7). The UMAP plot depicted 13 distinct cell clusters from the three respective cell samples (Fig. 5B). LymphH cells had the largest number of subclusters (7), with clusters 1-5 being very similar in composition (SI Fig. 7). Clusters 6 and 7 (lymphH6 and lymphH7) exhibited higher levels of pro-survival/cell cycle regulation genes (*BIRC5, TOP2A, CENPF*) associated with metabolically active cancer pathways. We observed the lung CD44^high^ (lungH) cells divided into three different clusters. The majority were localized to cluster lungH1 and expressed lower levels of metastatic cancer cell markers (*ALDH3A1, VIM, TCEAL9*). Similar metastatic cancer markers were upregulated in cluster lungH2. This subpopulation further displayed upregulated markers associated with the microenvironment and EMT (*AEBP1, ITGB5, FN1*).

### EMT and proliferation markers show distinct tissue-specific gene expression

We identified transcriptomic changes relevant to cancer metastasis that drove the designation of each cluster. Guided by the bulk RNA-seq results, we examined the expression of EMT and metastatic progression markers (Fig. 5C). We used dot plots to visualize the average expression of each gene and the percentage it was expressed in the sample set. As expected for progressive disease, we observed genes associated with primary tumors and low metastatic lesions (*CLDN3, KRT8, CDH1, CDH2*) primarily in the lymphH populations. Expression of these genes diminished at more distal sites (lung-derived samples). Conversely we observed genes associated with aggressive metastatic disease and EMT (*VIM, SPARC, ZEB1, SNAI2, CTSB*) upregulated in lung populations compared to lymph nodes. Changes in the expression of breast cancer marker CD63 were also observed during disease progression. Interestingly we observed similar gene expression changes for EMT in clusters lymphH6, lymphH7, and lungH3.

To further examine the changes in gene expression, we performed pseudo-time analysis of the single cells using Monocle2. The metastatic trajectory of the cells showed distinct clusters across the pseudo-time (Fig. 5E). We identified the composition of the cells by coloring the pseudo-time map with the tissue of origins (Fig. 5F). We observed that the majority of cells at the beginning of the pseudo-time (0) are lymphH-derived with a few lungL cells. Lung-derived cells became prominent further on the graph around pseudo-time 8. Interestingly, we observed lungL cells present across pseudo-time that cluster heavily towards the end of the graph (pseudo-time 12-16). We probed for changes in *VIM* and other EMT-relevant genes across pseudo-time and tissue of origin. As we previously observed in Fig. 5A, *VIM* expression increases over lymph node populations (Fig. 5G). *VIM* expression is highest in lung high populations and drops down in lung low populations. *CTSB*, a marker of invasion, showed relatively low expression across lymph nodes populations (Fig. 5H), but higher expression along pseudo-time in lungH and lungL populations.

Cellular proliferation markers were also analyzed via scRNA-seq (Fig. 5D). Increases in proliferative markers (*AURKA, MKI67*) were observed for clusters lymphH6 and lymphH7, with the greatest expression in lungH3. We also observed many similarities between clusters lymphH7 and lungH3 with regard to EMT- and proliferation-associated gene expression. These correlations could potentially signify the metastatic progression of the disease from the lymph node (lymphH7) to lung (lungH3). LungL clusters, by contrast, expressed lower levels of genes involved in proliferation. These clusters expressed higher levels of genes involved in cell cycle control and growth inhibition (*CDKN1A, CDKN1b, CCND3*). We further probed for changes in a panel of proliferation-associated genes across pseudo-time and tissue of origin using Monocle2 (SI Fig. 8).

### Cellular metabolism and extracellular remodeling markers show distinct tissue-specific changes

We identified gene expression markers relevant to cellular metabolism and extracellular remodeling that drove the formation of each cluster. For example, lungH3 cells expressed upregulated levels of genes associated with cell cycle and intracellular metabolism (*TOP2A, CENPF, HTRA1, STMN1*) (SI Fig. 7). These data suggested that cells within the lungH3 cluster exhibit the highest metastatic propensity of the lung subsets. We further observed an increase in insulin-like growth factors (*IGBFP4, IGBFP7*) across different lung-derived clusters, specifically in the lungL2 cluster (Fig. 6A). Interestingly, the expression of *PDHA1*, a critical component for pyruvate to acetyl-CoA conversion, was primarily upregulated in lungL2, lymphH7, lymphH6. We probed for changes in these metabolism-related genes across pseudo-time and tissue of origin using Monocle2. As we previously observed in Fig. 6A, *IGBFP7* expression was significantly upregulated in lungL single cell populations as pseudo-time progressed (Fig. 6B).

**Figure 6.**
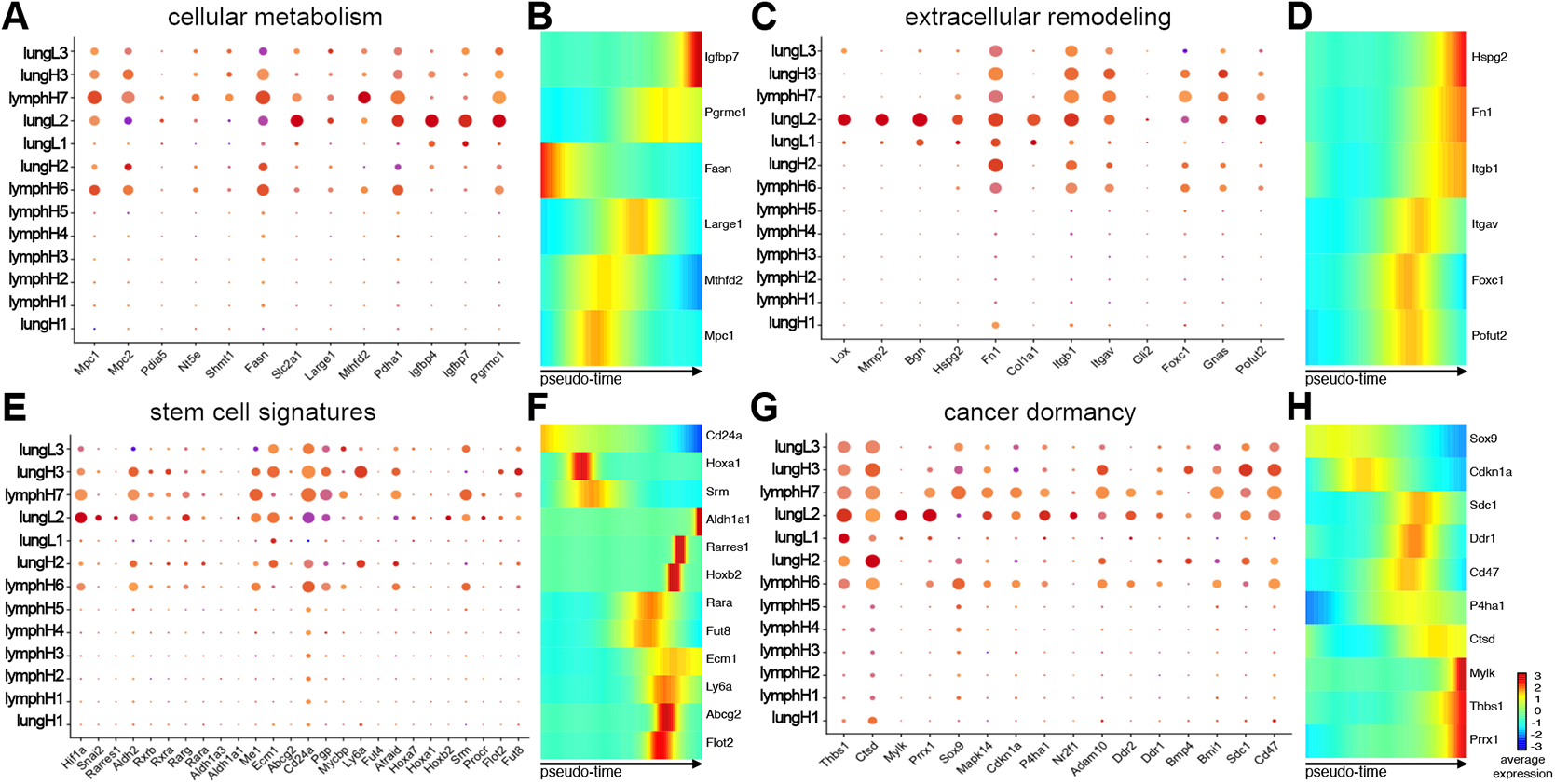
Single cell profiling revealed tissue-specific changes. **(A)** Dot plot analysis of **(A)** cancer metabolic markers relevant to cancer progression for select clusters guided by the bulk RNA seq data. **(B)** Heat maps of differentially expressed genes relevant to cancer metabolism were identified across the pseudo-time using Monocle2. The color gradient indicates the average expression across the pseudo-time, trending from dark blue to red. Similar dot plot analyses and pseudo-time heat maps are shown for markers relevant to **(C-D)** tumor microenvironment remodeling, **(E-F)** stem cell signatures, and **(G-H)** cell dormancy.

Guided by bulk RNA-seq results, we further examined extracellular remodeling markers using scRNA-seq (Fig. 6C and SI Fig. 7). We observed the most dynamic expression of extracellular remodeling-associated genes in the three lungL clusters. The lungL1 cluster had distinct differences in gene expression relevant to extracellular matrix interactions (*MGP, BGN, CCN2, FN1, ITGB1*). LungL2 cells expressed similar genes, along with upregulated genes associated with immunosuppressive proteins (*SLPI*) and embryonic glandular hormone (*PRL2C3*). However, cells in the lungL3 cluster lacked significant expression of extracellular remodeling genes that were present in lungL1 and lungL2. We investigated changes in extracellular remodeling-relevant genes (*FN1, ITGB1*) across pseudo-time and tissue of origin using Monocle2 (Fig. 6D). *FN1* had relatively low expression across lymph nodes populations, but increased expression was observed in lungH and lungL cells populations starting at pseudo-time 10. *ITGB1* was prominently expressed across most lungH, lungL, and some lymphH cell clusters (Fig. 6C). However, pseudo-time analysis of the single cells only attributed a significant upregulation of *ITGB1* expression in lungL cells at the end of the pseudo-time (Fig. 6D).

### CD44^low^ lung-derived cell lines harbor markers related to cancer stem cells and tumor dormancy

Although CD44 expression has traditionally been used as a marker of metastatic disease, recent publications have demonstrated that cellular expression of CD44 fluctuates during cancer progression.^(C. Yang et al., 2019)^ This has prompted other markers to be used in parallel with CD44 when examining metastatic potency and cancer-initiation capability of cells. Toward this end, we used scRNA-seq to probe for markers of CSC and tumor-initiation in the breast cancer model (Fig. 6E). We observed an increase in stem cell associated genes (*ALDH1A1, ECM1, ME1, ABCG2, and SNAI2*) with a subsequent decrease in differentiation marker, *CD24A*, in lungL1 and lungL2 clusters. Interestingly, lungL2 had the greatest upregulation in *HIF1A*, a known marker of hypoxia and poor prognosis in breast cancer. We previously observed that cells in lungL3 cluster separate from the other two lungL clusters (Fig. 5B). These cells have unique changes in gene expression relevant to embryonic and cancer stem cells (*KDEL1, PRL2C3, EBP1*, SI Fig. 7). Similar to clusters lungL2 and lungL3, stem cell genes *ALDH1A1, ABCG2*, and *SNAI2* were upregulated in lungL3 cells. However, other stem cell associated genes (*ME1, HIF1A*) were downregulated. Additionally lungL3 displayed an increase in *CD24A*, a marker of cellular proliferation and decreased stemness. Similar to the bulk RNA sequencing data, we saw an increase in *ALDH1A1* expression and stem cell survival metabolomic markers from the RA family in lungL2 and lungL3 clusters that were not present in lungL1 cells. In contrast, CD44^high^ clusters lungH3, lungH2, lymphH6, and lymphH7 exhibited an increase in genes regulating growth and invasion (*ECM1, CD24, FUT8*). While these CD44^high^ clusters showed a downregulation in stem cell markers *ALDH1A1* and *ABCG2*, they did show an increase in *ALDH2*, a stem cell marker less common in breast cancer.

We examined the changes in genes associated with cellular differentiation and stem cell capabilities (*CD24A, ALDH2*) using Monocle2 (Fig. 6F). *CD24A* expression was upregulated in lymph node populations, then downregulated as pseudo-time moved into the lung populations. Upregulation of stem cell marker *ALDH2* was observed across a select amount of cells for all cell types across pseudo-time (Fig. SI8)

Cancer dormancy is a formidable obstacle in breast cancer research and treatment. ^(Kim et al., 2012)^ Previous work has shown that some dormant breast cancer cells have similar genomic profiles as CSC.^(Hen & Barkan, 2020; Sosa, Bragado, & Aguirre-Ghiso, 2014)^ We examined whether our cell lines expressed markers of cancer dormancy and cancer stem cell-associated markers. Similarly to the bulk RNA-seq analyses, we saw an increase in genes regulating cell proliferation and increasing tumor dormancy in the lungL2, and in lungL1 to a lesser extent (Fig. 6G). This pattern was not sustained in the lungL3 subcluster. Instead, lungL3 showed a decrease in dormancy-associated genes *FN1, CD47*, and *THBS1*. Interestingly lungL3, lungL2 and lungH3 showed increased expression of breast cancer dormancy cell-associated maker *SDC1*.^(Gotte et al., 2006; Ibrahim et al., 2017)^ Master regulator of morphogenesis, *SOX9*,^(Grimm et al., 2020)^ was upregulated in lungL3, lymphH7, and lymphH6 subclusters. We further probed the changes in cancer dormancy-relevant genes *CTSD, THBS1* across pseudo-time and tissue of origin using Monocle2. *CTSD* expression fluctuated across lymph node populations (Fig. 6H). The expression of *CTSD* was upregulated through lungH populations and plateaued in the lungL populations across pseudo-time. *THBS1* exponentially upregulated expression at pseudo-time 10 in lung high and lung low populations (Fig. SI8).

## Discussion

Breast cancer comprises cancer cell subpopulations that are genetically and biologically different.^(Martelotto, Ng, Piscuoglio, Weigelt, & Reis-Filho, 2014)^ Although such intratumoral heterogeneity is critical to understanding the disease, traditional cancer cell lines models have difficulties recapitulating the complexity.^(Haynes, Sarma, Nangia-Makker, & Shekhar, 2017; Liu et al., 2018)^ In this study, we established a novel suite of organ-derived metastatic cell lines and subsequently performed a comprehensive transcriptome analysis of cancer progression. As previously shown in 4T1 cell lines^(Aslakson & Miller, 1992)^, models of metastatic breast cancer are invaluable for the advancement of the field. The MMTV-PyMT mouse model of breast cancer is the most popular transgenic preclinical system to study mammary tumor progression and metastatic disease translatable to patients.^(Le Voyer et al., 2000; Lifsted et al., 1998)^ Here, we successfully isolated primary cell lines from four different organ-derived tissues. The primary cancer cell lines were isolated based on their expression of CD44^low^/EpCAM^high^ or CD44^high^/EpCAM^high^ signatures.

The suite of MMTV-PyMT derived cell lines was subjected to comprehensive transcriptome analysis using bulk RNA-seq and scRNA-seq. Although there were shared GO terms across all cell lines belonging to either CD44 signature, we identified numerous changes in gene expression that were organ-specific and promoted metastatic homing. ^(W. Chen et al., 2018)^ Changes in gene expression related to EMT and the metastatic cascade were also identified with similar expression across both transcriptome pathways. As expected distal metastatic-derived tissues such as the lung and BM had more differential genes expressed than lymph node-derived cell lines compared to the primary tumor.

Using scRNA-seq on select organ-derived metastatic cell lines, we found that the cell lines exhibited a range of intratumoral heterogeneity. Although most MMTV-PyMT publications have focused on metastasis from the primary tumor to the lungs, we identified additional distal metastatic cells in the BM. The BM-derived cell lines were characterized and showed an increase in previously published markers such as *RANKL, OPN* (*SPP1*), and *IL2*.^(Hanahan & Weinberg, 2011)^ Further studies are necessary to better characterize these cells and obtain cleaner sequencing analyses. Specifically, it would be prudent to examine the differential gene expression of this model compared to other well-established bone-marrow metastatic models of tumor latency and cancer dormancy.ref Collectively, these transcriptome changes could contribute to sub-clonal evolution during cancer progression across the different metastatic niches.

Although CD44 has traditionally been used to monitor cancer metastasis, recent studies from Gao and corkers showed that its expression fluctuates during metastatic progression.^(C. Yang et al., 2019)^ We observed similar changes in CD44 expression from initial isolates compared to cells used in subsequent transcriptome analysis. Furthermore, recent studies with MDA-MB-231 breast cancer models showed that CD44^low^ cells possessed stem cell like properties that CD44^high^ lacked.^(K. Chen & Fraley, 2020; Y. Li & Laterra, 2012)^ CD44^low^ populations could regenerate both more CD44^low^ and CD44^high^ cell clusters, where CD44^high^ could only replicate themselves. We observed a similar increase in stem cell markers in CD44^low^ but not CD44^high^ cells. These contradictory results suggest that more research is needed to better understand and characterize CD44 expression in breast cancer.

Although CD44 has been correlated to metastatic progression and CSC, the fluctuating expression levels of this marker complicates its use as a sole classifier of cellular phenotypes. There are thus many combinations of markers currently used to isolate CSC, with no one set being universally accepted.^(K. Chen & Fraley, 2020; Y. Li & Laterra, 2012; Nowell, 1976; Sachidanandam et al., 2001)^ We examined a handful of CSC markers commonly associated with breast cancer (*ALDH1A1, ABCG2, PGP, FUT4*). CSC associated markers were found in the both bulk RNA-seq and scRNA-seq data. The expression of the CSC-associated markers was highest in lung-derived CD44^low^ cells compared to CD44^high^ cells. Upregulation of other CSC-associated markers, including the stem cell survival signaling member WNT, was also observed in the CD44low cells, corroborating our hypothesis that these cells possess stem-like properties.^(Ma et al., 2012)^ scRNA-seq further revealed three distinct clusters for lung-derived CD44^low^ cells on the UMAP. Unlike the lung-derived CD44^high^ groups, lungL1 and lungL2 retained populations of stem cells but dramatically increased their expression of mesenchymal-like signaling pathways involved in cancer cell maintenance and dormancy (*TSP, TNC, BMP*) ^(De Angelis et al., 2019; Ghajar et al., 2013; Montagner & Sahai, 2020)^. Interestingly, lungL3 was a much smaller, separate cluster that lacked the dormancy markers and instead expressed high levels of stem cell markers. The diversity of CD44^low^ clusters could be explained by heterogeneity within the different stages of metastatic progression and along the EMT spectrum. Another possibility is that lungL1 and lungL2 clusters might be cancer associated fibroblasts while the lungL3 might be a distinct cluster of CSC cells.

Breast cancer progression relies on CSC, to initiate tumor cell growth during metastatic dissemination and colonization of new organs. Previous work from Weinberg has shown that TICs originating in the luminal cell layer of the mammary gland rely on EMT initiating transcription factors (TFs) for cellular dedifferentiation.^(Ye et al., 2015)^ These factors activate signaling pathways distinct from TICs originating from the basal mammary gland. Bulk RNA-seq data allowed us to further characterize the lung-derived cell lines based on expression of EMT-TFs known to induce TICs. For example, increased expression of the EMT-TF Snail was observed in lung CD44^low^ cells compared to CD44^high^ cells. Although both CD44 signatures of lung-derived cells expressed the EMT-TF binding promoter and TIC master regulator *ZEB1*, lung-derived CD44^low^ cells also showed a greater increase in the basal cell associated EMT-TF Slug as compared to CD44^high^ cells. Weinberg and others have shown that Slug and Snail both bind and regulate *ZEB1* expression ^(Ye et al., 2015)^;however, the expression of *SLUG* is associated breast cancers that arise from cells containing normal mammary epithelial stem cells in the basal compartment *SLUG* expression is also associated with highly dedifferentiated breast cancer cells found in the advanced and final stages of metastatic disease. Here, we harvested MMTV-PyMT mice well after the time period sampled by Weinberg. The most striking result from the lung-derived cells was that the stem cell associated features of tumor-initiating cells were observed most in CD44^low^ opposed to CD44^high^ expressing cells that have previously been correlated to CSC.

Further work examining the effects of the microenvironment as a key regulator of cellular plasticity adds to the theory that “dedifferentiated” non-CSCs can undergo processes that endow them with CSCs-like properties. ^(Y. Li & Laterra, 2012; Takebe et al., 2011)^ These complex studies highlight the need for further investigation into the possible origins of CSCs in relations to the surrounding microenvironment. Modification of cellular metabolism along with remodeling of the tumor microenvironment can be achieved through the collective change of different groups of genes. The modulation of certain genes have been shown to subsequently facilitate cancer stem cells phenotypes such as TIC that lead to metastatic colonization. Based on the gene expression changes dictating cellular metabolism and matrix remodeling, we hypothesized that lung low cells could harbor some cancer stem cell properties. Expression of some cancer stem cell markers did identify expression of relevant genes in this set of cells.

To better understand the origins of the CD44 signatures, we examined the scRNA-seq results as they projected over pseudo-time using Monocole2. From the three different cell lines, we identified that lymph node-derived CD44^high^ cells were projected to give rise to lung-derived cells of both signatures. Based on the metastatic cascade of organ tropism, the proximal location of the lymph nodes to the primary tumor is well documented to be the initial site of metastatic lesions before advancing to the more distal lungs. Future studies would benefit from expanding the analysis to include lymph node-derived CD44^low^ cell lines to determine how the signature affects the projection of the tumor initiating cell population.

Collectively, we established organ-derived cancer cell lines from different metastatic niches. Comprehensive transcriptomic analysis was performed and revealed the impacts of heterogeneity on cancer progression. These data will improve our understanding of the metastatic cascade and tumor heterogeneity in breast cancer will dramatically improve targets for therapies.

## Supporting information

SI Fil 4a

## Acknowledgements

This work was supported by the U.S. National Institutes of Health (R01 GM107630 to J. Prescher), and a UCI Opportunity Center for Complex Biological Systems Grant. A. Ionkina was supported by an institutional Cancer Biology Training Grant (T32-CA009054) and G. Balderrama-Gutierrez was supported by a UC Mexus-CONACYT fellowship. We thank members of the Prescher, Mortazavi and Kessenbrock laboratories for helpful discussions.

## Supporting Information

### Methods

#### Mammalian cell culture

Unless otherwise stated, cell lines were cultured in DMEM (Corning) supplemented with 10% (vol/vol) fetal bovine serum (FBS, Life Technologies), penicillin (100 U/mL), and streptomycin (100 µg/mL). Cells were maintained in a 5% CO_2_ water-saturated incubator at 37 °C.

#### MMTV-PyMT metastatic cell lines as models of breast cancer

Mouse experiments were approved by the UC Irvine Animal Care and Use Committee. Tumor bearing organs were harvested from 10-12 week females FVB/NJ MMTV-PyMT mice (courtesy of the Kessenbrock laboratory, UCI). Samples were processed mechanically and chemically to dissociate tissues into single cell suspensions as previously published (Lawson et al., 2015). Primary single cell suspensions were enriched for cancer cells over the course of 1 month in vitro incubation by exploiting differences in cellular nutrient requirements and growth differentials. During this time course, primary single cell suspensions were enriched for cancer cells by culturing cell lines in 5% FBS. Cultures were selected for immortalized cancer cells in vitro by passaging the flasks 3 times a week. Differences in cellular adhesion properties between fibroblasts and epithelial cancer cells were also exploited in vitro through 3 mins versus 7 mins incubations with trypsin (Schor et al., 1979). The month long process resulted in the enrichment of cancer cell lines in the surviving in vitro cultures prior to FACS sorting.

Primary cell lines were processed for FACS sorting (Institute for Immunology Flow Cytometry Core, UCI) as previously reported (Liao, Makris, & Luo, 2016). Cancer cells were isolated by EpCAM (BioLegend 118213) and CD44 (BioLegend 103027) cell surface expression levels (Marhaba et al., 2008); isolated cancer cells expressed either CD44^low^/EpCAM^high^ or CD44^high^/EpCAM^high^ cell surface markers. Antibody labeling was performed using manufacture’s protocols (BioLegend, USA). Previously sorted MMTV-PyMT MFP-eGFP cell lines (VO), (courtesy of the Kessenbrock laboratory and Lawson laboratory, UCI) were used as positive controls. Fibroblast cell lines (3T3 and MMTV-PyMT-derived fibroblasts, isolated during culturing process above) were used as negative controls during sorting.

#### Primary cell line metastatic propensity validation in vivo

MFP-derived cells (100,000 cells/injection) were injected bilaterally under the fourth gland of disease free, 4-week old FVB/NJ female mice. Control VO-eGFP luciferase-expressing cells were injected as a control to monitor the estimated tumor growth. Palpable primary tumors were detected in all mice within 3-4 weeks post injection. All animals developed primary tumors. Metastatic cell populations were identified by harvesting and processing the organs as described above via FACS analysis. Cancer cells were isolated using cell surface expression of CD44 and EpCAM. The experiment was performed over 4 different biological replicates.

#### PCR analysis

gDNA was isolated from all MMTV-PyMT cell lines and control samples using Zymo (California, USA) quick-DNA miniprep kit (Cat #: 11-317AC). Ear clippings from PyMT positive male and female mice (courtesy of the Kessenbrock laboratory, UCI) were used as positive controls. gDNA samples isolated from 4T1 (ATCC CRL-2539) cell lines were used as negative controls. PCR amplification conditions and PyMT antigen detection were completed using the standard Jackson Labs genotyping protocol (Guy, Cardiff, and Muller 1992).

#### Bulk-RNA-seq

For each tissue derived cell line, total RNA was extracted using the QIAGEN RNeasy kit with 2 replicates per sample. A modified SMART-seq2 protocol was used to generate cDNA and Nextera XT DNA Sample Prep Kit to build Illumina libraries. Samples were sequenced on a NextSeq500 with a min depth of 10 M reads. Raw reads were aligned to the mm10 genome with STAR(Dobin et al., 2013) and quantification was performed using the GENCODE v21 annotation of the mouse genome using RSEM (Li & Dewey, 2011). Count matrices for differential expression analysis were used as input for EdgeR. An Exact test was used for calling differential expressed genes with logFC > 2 and a p value < 0.05. EnrichR and metascape were used for Gene ontology analysis.

#### Single-cell RNA-seq methods

Cell lines that originated from Lung with CD44^high/low^ signatures were identified as the samples with most transcriptional changes and were selected for single-cell analysis, along with a Lymph node high sample. Single-cell suspensions from these tissues were used as input for the ddSeq platform and cDNA synthesis as well as library prep was done following the SureCell™ Whole Transcriptome Analysis 3’ Library Prep Kit. The bioinformatic pipeline included Ddseeker (Robinson, McCarthy, & Smyth, 2010), a custom demultiplexing script to generate individual fastq, while kallisto (Bray, Pimentel, Melsted, & Pachter, 2016) was used to quantify the transcripts in our sample using the mm10 and annotation GENCODE v21. Single-cell analysis was done using Seurat v3.2.3 (Bray et al., 2016). Cells with more than 250 genes and less than 10% mitochondrial reads were used for the analysis. Monocle2 (Bray et al., 2016) was used to infer pseudotime progression. Min. read depth 34 M.

## Data availability

Fastq files for bulk and single-cell datasets as well as their corresponding processed matrices are available in GEO (Accession number: GSE165393)

**SI Figure 1.**
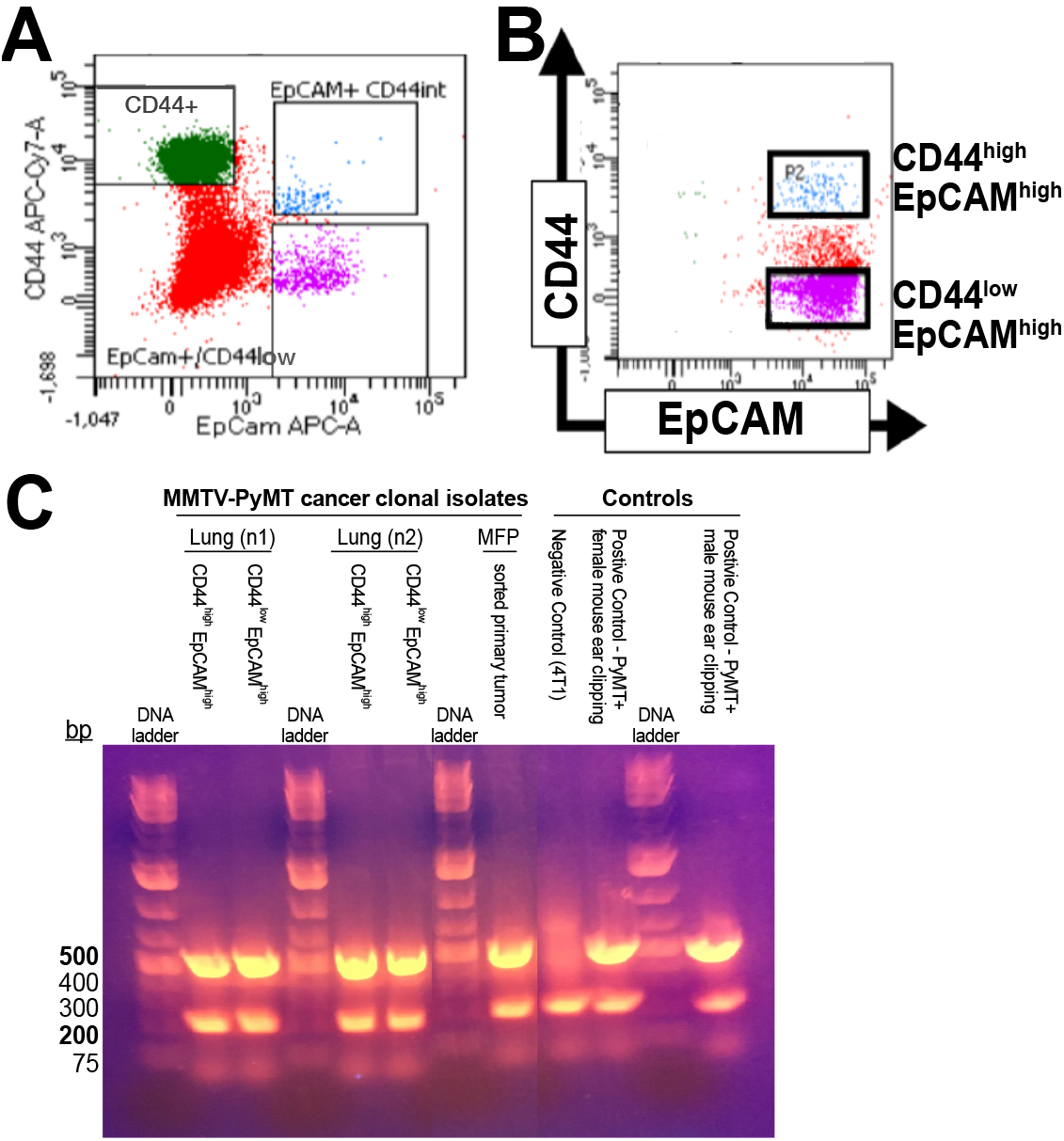
Metastatic cell line derivation and validation. **(A)** Representative flow cytometry plot for metastatic tissue-derived isolates prior to sorting. The desired CD44^low^/EpCAM^high^ and CD44^high^/EpCAM^high^ cell populations and associated gates are highlighted. These cells were sorted and for transcriptomic analysis. **(B)** Clonal isolates from **(A)** recapitulate metastatic disease in vivo. Sorted cells were reimplanted into disease-free FVB mice. Once tumors formed, distal organs were collected and examined for metastatic lesions. A representative FACS plot for metastatic cells harvested from lymph node tissue upon tumor reimplantation in FVB mice. The expected CD44^low^/EpCAM^high^ and CD44^high^/EpCAM^high^ populations are highlighted. **(C)** PCR was used to confirm the presence of PyMT viral antigen using gDNA extract from cell lines and controls. Samples with amplified 500bp and 200bp bands are positive for PyMT antigen expression.

**SI Figure 2.**
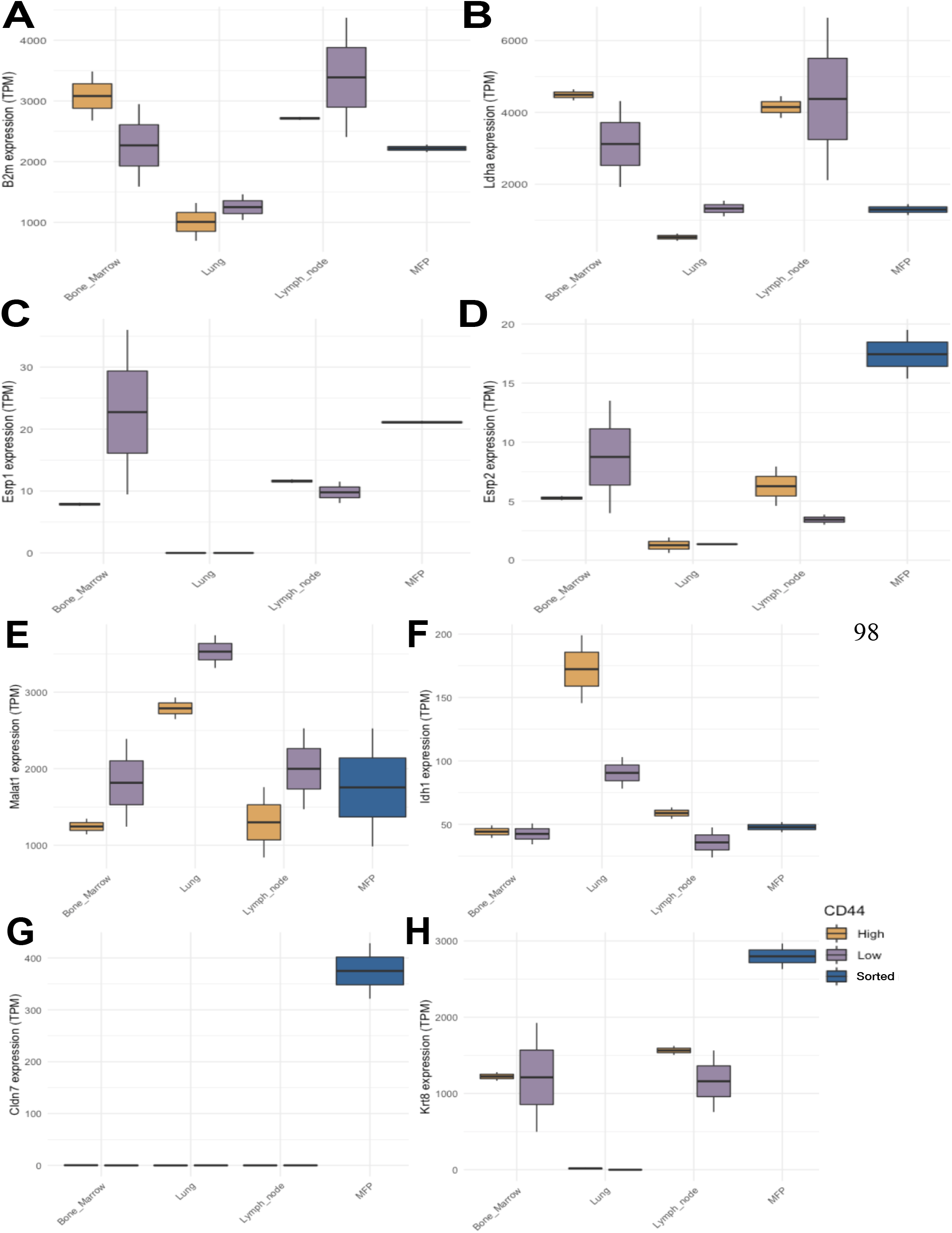
Established breast cancer transcripts identified in MMTV-PyMT cell lines. Bulk RNA sequencing analysis of bone marrow, lung, and lymph node-derived cell lines was performed and transcript levels for a panel of breast cancer markers were measured. For (A-H), expression of gene transcripts (TPM, transcript per million) for CD44 high expressing cells (yellow), CD44 low expressing cells (purple), and MFP CD44 expressing cells (blue) are shown.

**SI Figure 3.**
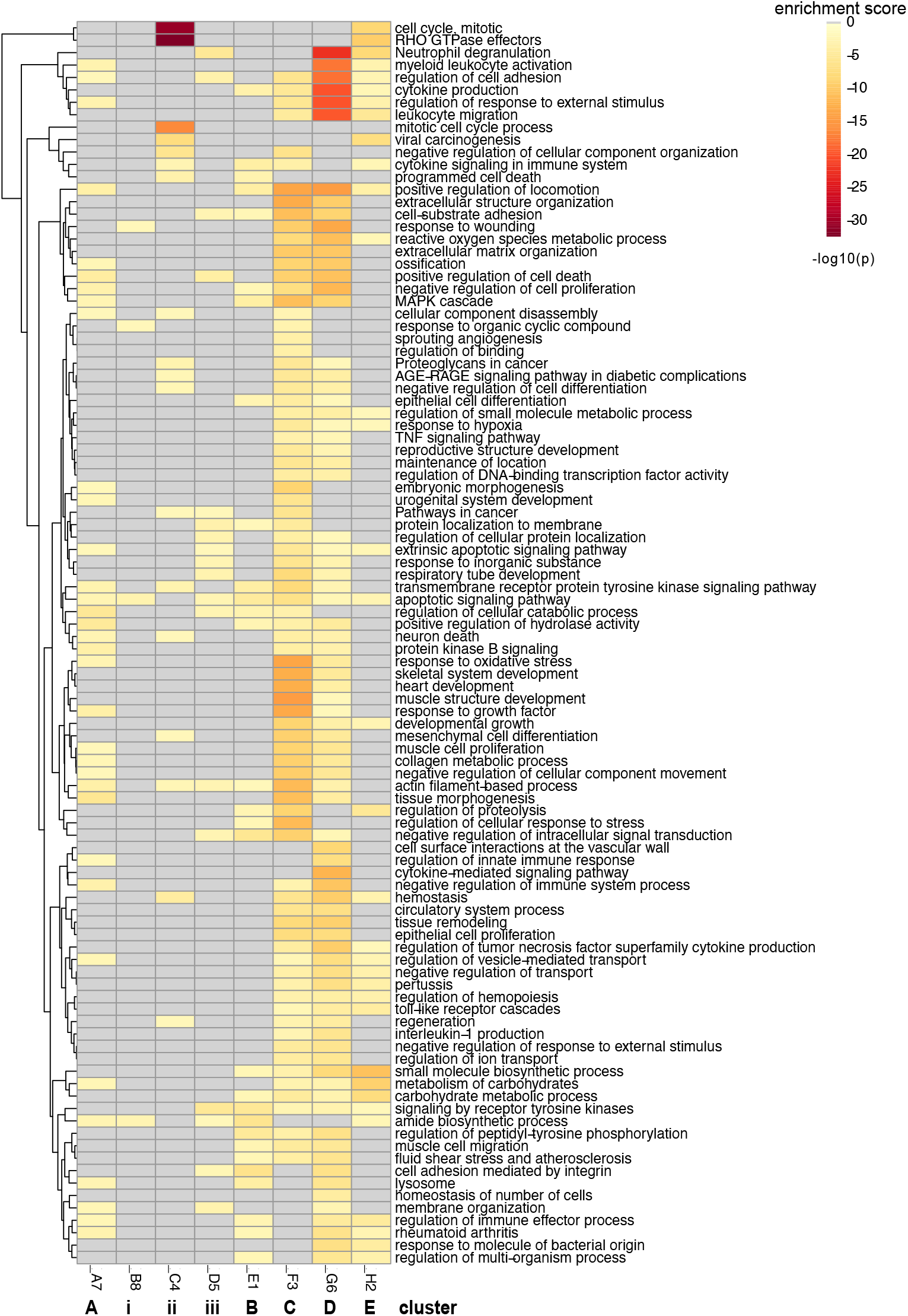
Complete list of enriched GO-terms and pathway analysis from Fig 1D. Heat map shows the enrichment scores of the pathways for each cluster (A-E; i-iii).

**SI Figure 4.**
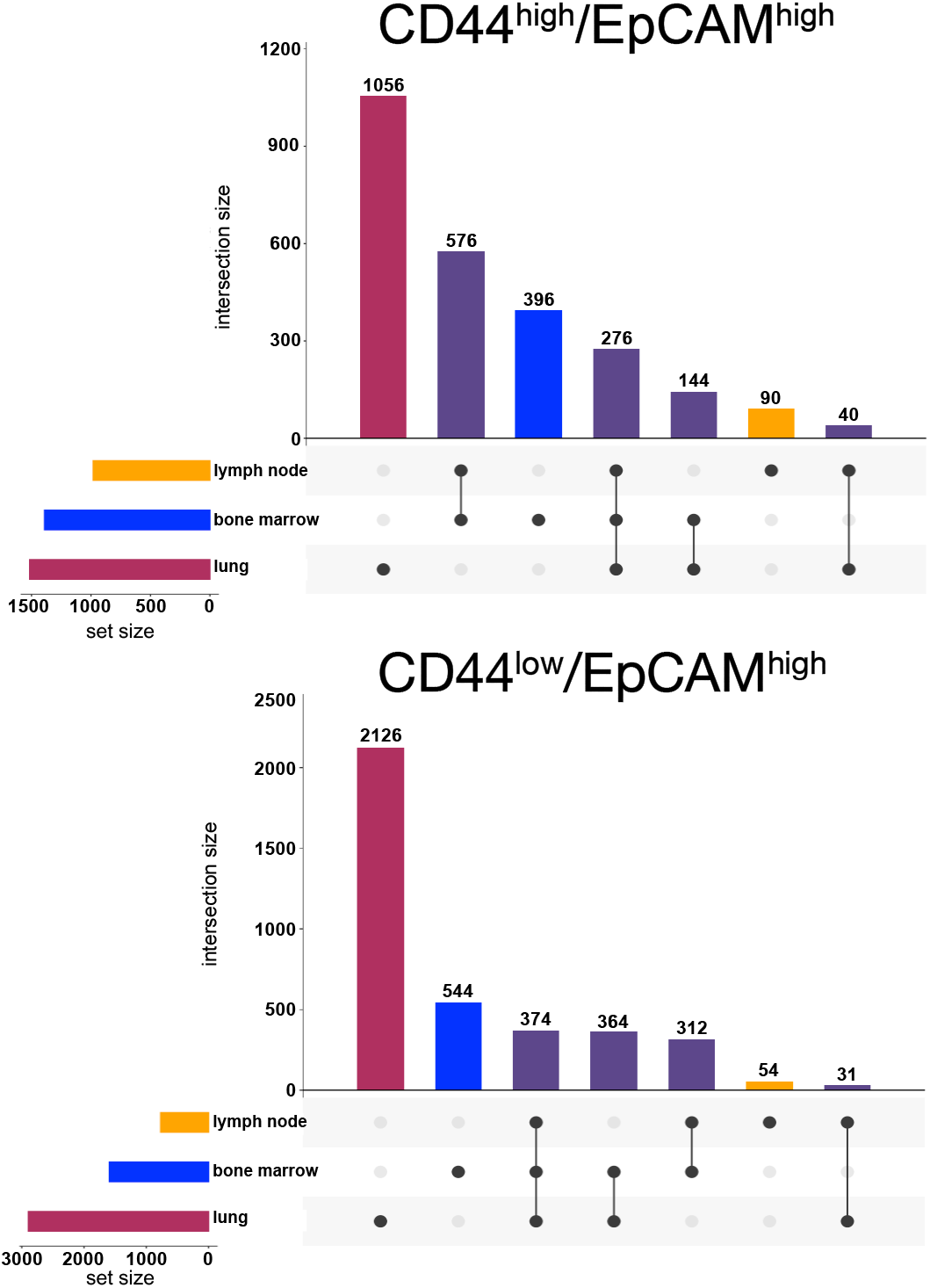
Box and whisker plot comparing the number of upregulated genes in CD44^high^/EpCAM^high^ (top plot) and CD44^low^/EpCAM^high^ (bottom plot) expressing cells from the different metastatic tissues of origin. Distinct gene expression signatures are shown with single black dots corresponding to a specific tissue-derived clonal isolate. Genes that are shared between different metastatic sites are represented by black lines that connect the specific samples. The total number of upregulated genes are denoted above each bar on the graph.

**SI Figure 5.**
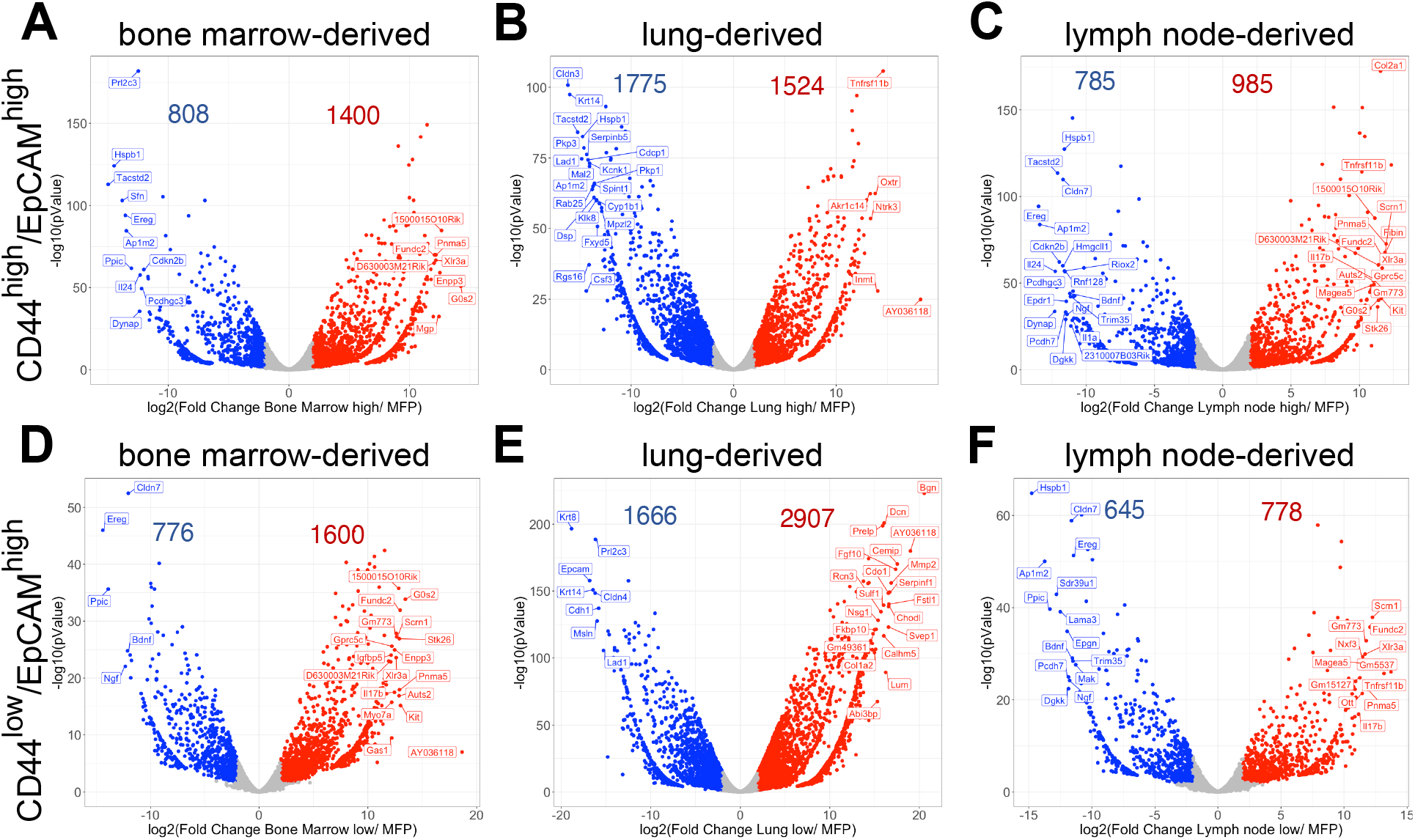
Volcano plots of DEGs from tissue-derived metastatic cell lines. **(A-C)** Volcano plots of DEGs from tissue-derived metastatic cell lines with CD44^high^/EpCAM^high^ expression compared to primary tumor samples. **(A)** Upregulated genes in bone marrow-derived (BM) isolates with CD44^high^/EpCAM^high^ expression are shown in red (1,400 genes), while 808 genes (blue) were upregulated in the primary tumor. **(B)** Upregulated genes in lung-derived isolates with CD44^high^/EpCAM^high^ expression are shown in red (1,524 genes), while 1,775 genes (blue) were upregulated in the primary tumor. **(C)** Upregulated genes in lymph node-derived (LN) isolates with CD44^high^/EpCAM^high^ expression are shown in red (985 genes), while 785 genes (blue) were upregulated in the primary tumor. **(D-F)** Volcano plots of differentially expressed genes from tissue-derived metastatic cell lines with CD44^low^/EpCAM^high^ expression when compared to primary tumor samples. **(D)** Upregulated genes in bone marrow-derived (BM) isolates with CD44^low^/EpCAM^high^ expression are shown in red (1,600 genes), while 776 genes (blue) were upregulated in the primary tumor. **(E)** Upregulated genes in lung-derived isolates with CD44^low^/EpCAM^high^ expression are shown in red (2,907 genes), while 1666 genes (blue) were upregulated in the primary tumor. **(F)** Upregulated genes in lymph node-derived (LN) isolates with CD44^low^/EpCAM^high^ expression are shown in red (778 genes), while 645 genes (blue) were upregulated in the primary tumor. The fewer differentially expressed genes amongst the tissue-derived isolates compared to the primary tumor is suggestive of the order in the metastatic cascade with LN-derived isolates bring the first metastatic site.

**SI Figure 6.**
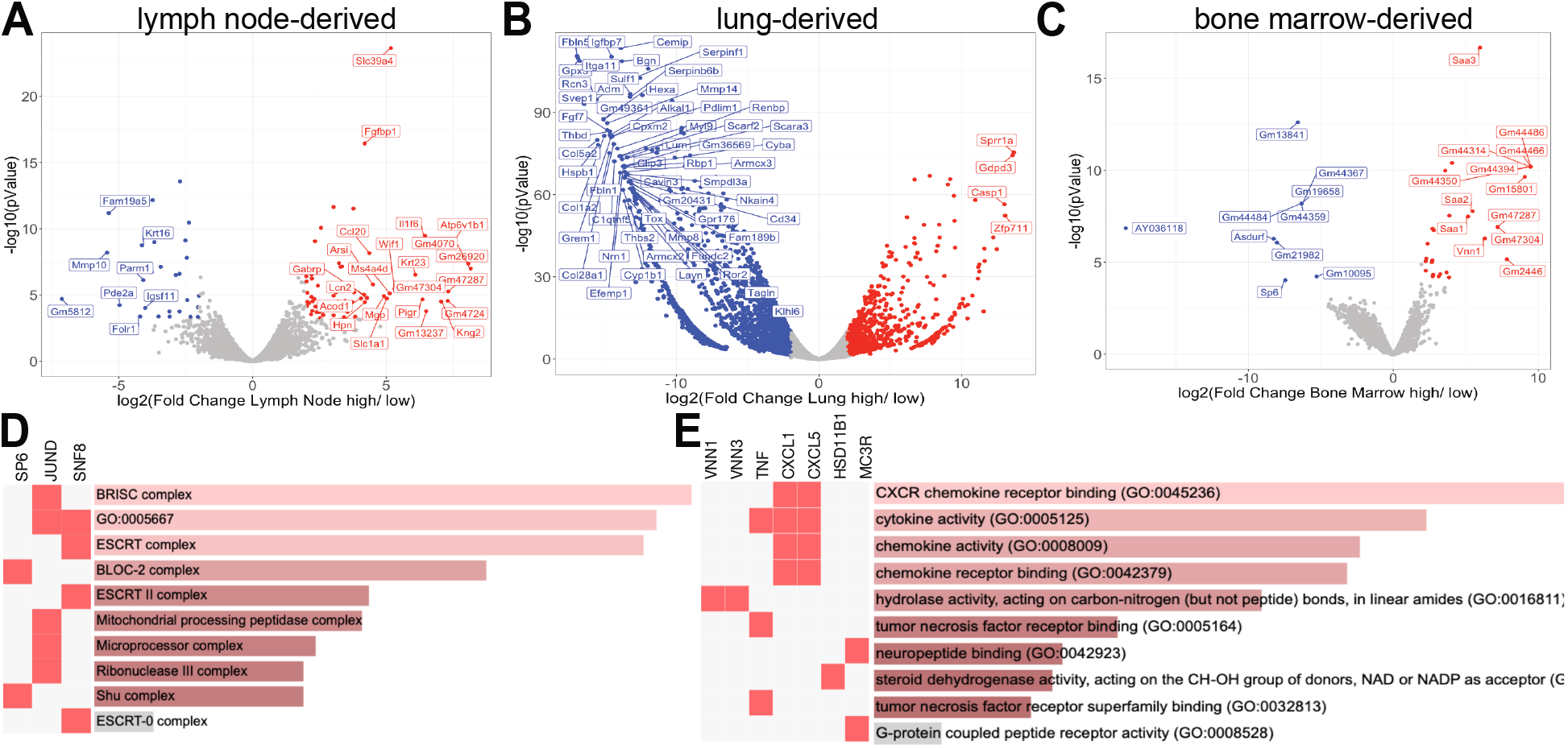
Analysis of DEGs from metastatic isolates based on CD44 expression. **(A**) Volcano plots showing DEGs in CD44^high^/EpCAM^high^ (red) versus CD44^low^/EpCAM^high^ (blue) from **(A)** lymph node-derived, **(B)** lung-derived, and **(C)** bone marrow-derived metastatic cells. **(D)** GO terms and relevant genes upregulated in CD44^high^/EpCAM^high^ bone marrow-derived cells. **(E)** GO terms and relevant genes upregulated in CD44^low^/EpCAM^high^ bone marrow-derived cells.

**SI Figure 7.**
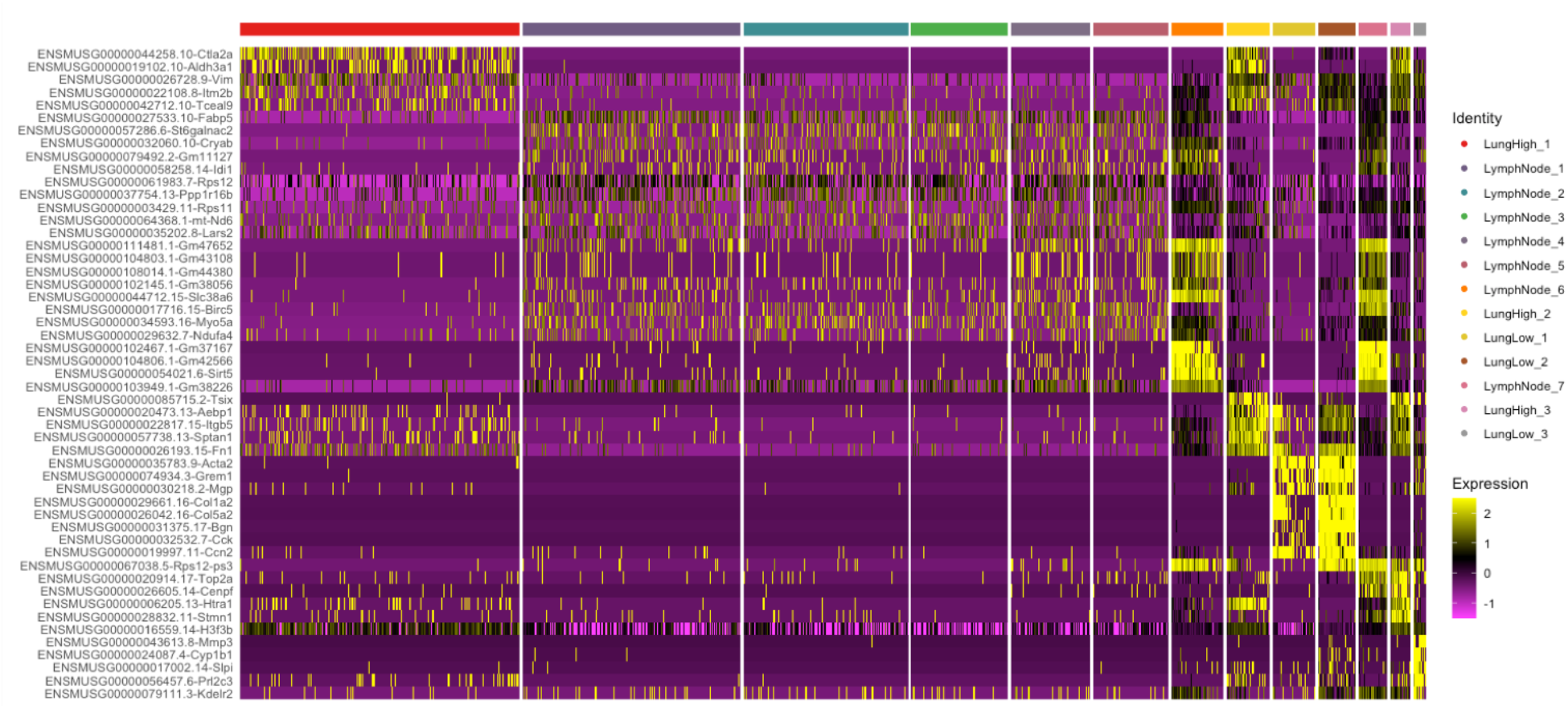
Hierarchical clustering analysis from single cell sequencing analyses. Single cell sequencing heat map showing 13 tissue-specific clusters.

**SI Figure 8.**
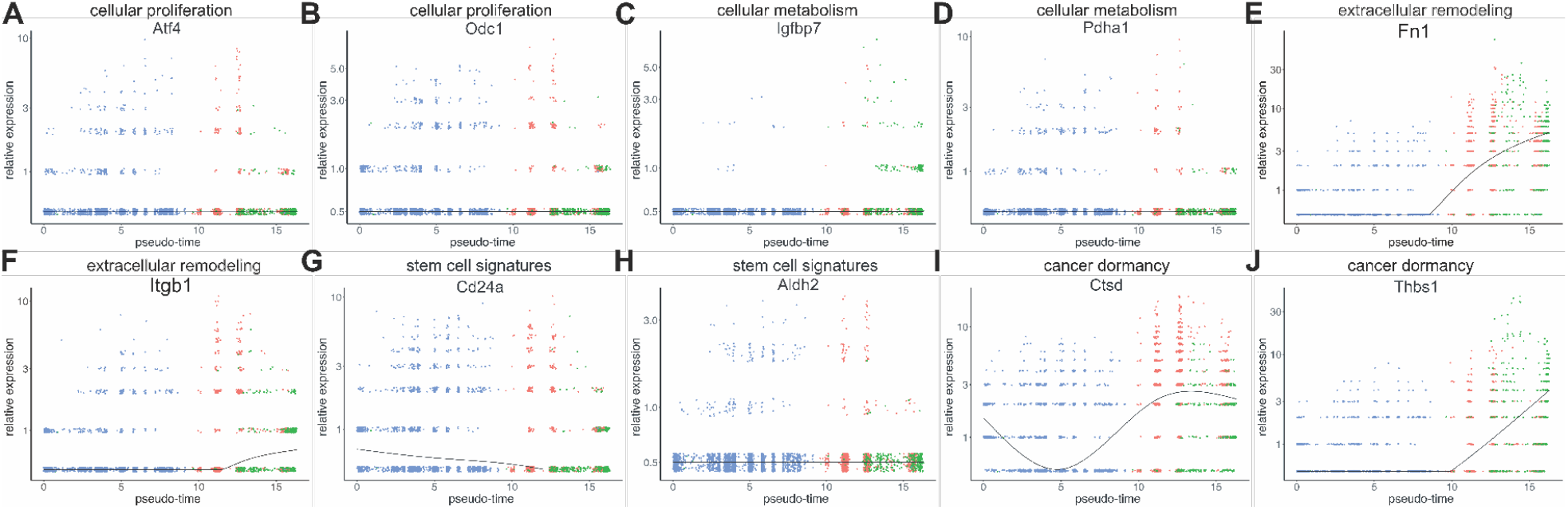
Monocle2 pseudo-time analysis of single cells. The metastatic trajectory of distinct cells clusters is shown. The composition of the cells was identified by coloring the pseudo-time map with the tissue of origins (as in Fig. 5A). Expression of cellular proliferation markers **(A, B)**, metabolism markers (**C, D)**, extracellular remodeling markers **(E**,**F)**, stem cell signatures **(G**,**H)**, and cancer dormancy markers **(I**,**J)** of single cells across pseudo-time. Trend line on graphs tracks the statistical significance of gene expression as it changes across pseudo-time.

**SI File 1. Allmetascape**

*File attached separately*

Complete list of enriched GO-terms and genes from the enriched pathway analysis from Fig 1D.

**SI File 2. Metastatic tissues derived vs MFP cell lines**.

*File attached separately*

Differential gene analysis comparing metastatic tissue derived-cell lines (LU, BM, and LN) with CD44^low^ and CD44^high^ expression against the MFP-derived cell line (primary tumor)

**SI File 3. lowvshigh**

*File attached separately*

Differentially expressed gene analysis comparing CD44^low^ and CD44^high^ samples derived from each metastatic tissue.

**SI File 4. Genes and GO terms for Fig 3A**

*File attached separately*

Complete list of enriched GO-terms and genes from the enriched pathway analysis from all upregulated genes from CD44^low^ and CD44^high^ samples across all tissues (from Fig 3A).

**SI File 5. Genes and GO terms for Fig 3B**

*File attached separately*

Complete list of enriched GO-terms and genes from the enriched pathway analysis from all upregulated genes from CD44^low^ and CD44^high^ samples across lymph node-derived cell lines (from Fig 3B).

**SI File 6. Genes and GO terms for Fig 3C**

*File attached separately*

Complete list of enriched GO-terms and genes from the enriched pathway analysis from all upregulated genes from CD44^low^ and CD44^high^ low samples across lung-derived cell lines (from Fig 3C).

